# Disrupted Hippocampal Theta-Gamma Coupling and Spike-Field Coherence Following Experimental Traumatic Brain Injury

**DOI:** 10.1101/2024.05.30.596704

**Authors:** Christopher D. Adam, Ehsan Mirzakhalili, Kimberly G. Gagnon, Carlo Cottone, John D. Arena, Alexandra V. Ulyanova, Victoria E. Johnson, John A. Wolf

## Abstract

Traumatic brain injury (TBI) often results in persistent learning and memory deficits, likely due to disrupted hippocampal circuitry underlying these processes. Precise temporal control of hippocampal neuronal activity is thought to be important for memory encoding and retrieval and is supported by oscillations that dynamically organize single unit firing. Using high-density laminar electrophysiology, we found a loss of oscillatory power across CA1 lamina, with a profound, layer-specific reduction in theta-gamma phase amplitude coupling in injured rats. Interneurons from injured animals were less strongly entrained to theta and gamma oscillations, but both interneurons and pyramidal cells from injured animals became more strongly entrained to theta during periods of high theta power. During quiet immobility, sharp-wave ripple amplitudes were lower in injured animals compared to shams. These results reveal physiological deficits across brain states that may contribute to TBI-associated learning and memory impairments and elucidate potential targets for future neuromodulation therapies.

## INTRODUCTION

Cognitive deficits, including learning and memory impairments, are one of the most common consequences of TBI^1–3^. These deficits occur when complex, heterogeneous TBI pathologies affect circuits that support these processes. Local circuits within the hippocampus are especially important for learning and memory, as are interactions between the hippocampus and connected brain regions. Because these networks are dispersed, it is not surprising that TBI-associated learning and memory deficits have been reported both in the presence and absence of hippocampal pathology^4–7^. Neuronal pathology in hippocampal afferents such as the medial septum and para-hippocampal cortical regions can affect hippocampal processing, as can axonal damage along connecting pathways^8–12^. Additionally, other coalescing pathological changes such as neurotransmitter dysregulation^13–15^, inflammation^16,17^, blood brain barrier breakdown^18,19^, and metabolic dysfunction^20–23^, as well as changes in intrinsic neuronal and circuit excitability and synaptic plasticity^16,24–34^ are all thought to contribute to hippocampal dysfunction following TBI. However, it is unclear how these pathological changes collectively affect information processing in the intact hippocampus. Studying this process in awake subjects is important since network level physiological processes in the hippocampus have been shown to support learning and memory functions, and these electrophysiological processes can be predictive of behavior^35–39^.

Despite our understanding that hippocampal neuronal activity underlies many aspects of learning and memory, surprisingly few studies have investigated how TBI affects these processes in awake animals. Prior studies have reported a loss of oscillatory power in CA1 or broad changes in the firing properties of pyramidal cells post injury^5,40–45^. A recent study provided evidence that spike timing relative to theta oscillations, a process called spike-field coherence or entrainment, may be disrupted following TBI^46^. However, the cell type specificity of this change and the potential network mechanisms driving it remain poorly characterized. Spike-field coherence is important as hippocampal oscillations dynamically organize the firing of populations of neurons into cell ensembles, and the precise temporal control of cell ensemble firing is important for plasticity mechanisms underlying learning and memory^47–50^. Hippocampal oscillations coordinate ensembles in a complex manner, nesting high frequency gamma oscillations within a dominant lower frequency theta oscillation during active exploration^51,52^. Theta oscillations are important for synchronizing neuronal activity both within and across brain structures^53,54^, while gamma oscillations are thought to reflect local processing within a brain region and are supported by local interneurons^55^. Importantly the amplitude of gamma oscillations changes as a function of theta phase, a process known as theta-gamma phase amplitude coupling (PAC), which provides a mechanism whereby local processing across distributed networks can be temporally coordinated on a theta timescale^56–61^. Appropriate PAC and single unit entrainment to hippocampal oscillations are thought to be essential for cognition and have been directly linked to memory performance in humans^61–63^. Additionally, higher frequency sharp-wave ripples (SWRs) are known to support working memory processes and memory consolidation^64–68^, however, the effects of TBI on these processes have not been investigated.

We therefore utilized high-density laminar electrodes to characterize layer-specific pathophysiological changes to hippocampal oscillations and single unit activity from freely moving rats following the well-characterized lateral fluid percussion injury (_L_FPI) model of TBI. We found that, compared to controls, injured rats had layer-specific reductions in oscillatory power and PAC as well as decreased ripple amplitudes in CA1 which suggests a desynchronization of the hippocampal network across brain states. Additionally, we found that interneurons from injured rats are less strongly entrained to local theta and gamma oscillations, and that cell-type specific changes in entrainment and PAC are correlated with theta amplitude in injured animals. These results reveal disruptions to hippocampal network activity presumed to be fundamental to cognitive processing, which may contribute to TBI-associated learning and memory deficits. The theta amplitude dependence of these physiological changes may inform deeper mechanistic studies investigating TBI-associated cognitive deficits, and suggest physiological targets for future neuromodulation therapies.

## RESULTS

### _L_FPI decreases theta and gamma power across CA1 lamina

Previous studies have reported reductions in oscillatory power in the CA1 region of the hippocampus in awake animals following TBI^40,42,45^, but recordings from these studies were either specific to the pyramidal cell layer, or were obtained non-discriminately in the CA1 region. Layer-specific input combined with anatomically constrained inhibition from extensive interneuronal networks in CA1 create stereotyped oscillatory signatures that differ throughout the layers of CA1. Thus, TBI-associated changes observed in a specific layer could implicate specific inputs or interneuron subtypes targeting that layer. In order to test whether TBI disrupts network oscillations non-discriminately or in a layer specific manner, laminar recordings were obtained from the CA1 region of the hippocampus in freely moving sham (n=4) and injured (n=5) rats. Animals were subjected to _L_FPI (1.68-1.78 atm) or sham surgery then chronically implanted with multi-shank, high-density laminar electrodes in the CA1 region of the hippocampus ipsilateral to the injury/sham site. Recordings were obtained on three separate days within 6-11 days post-injury. On the first day of recording rats freely explored a familiar environment, followed by a novel environment. On the subsequent two recording days, electrodes were slowly advanced, then recordings were obtained in the familiar environment.

The use of laminar electrodes allowed us to localize electrode contacts across different CA1 lamina specifically *stratum oriens*, *pyramidale* (*st. pyr*), and *radiatum* (*st. rad*). To maximize single unit yields, most electrodes were localized to *st. oriens* and *st. pyr* on the first day. Thus, driving on subsequent days allowed for recordings to be obtained from *st. rad*. For each recording, electrodes from individual shanks were independently assessed to determine their precise location within the CA1 lamina. The *st. pyr* channel was defined as the channel with maximum power in the ripple frequency range (100-250 Hz; **Fig 1A**). As expected, individual pyramidal cells were localized around this channel (**Fig 1B**). The *st. rad* channel was defined as the local maximum of the current source density (CSD) sink from the mean SWR waveform (**Fig 1C**; local maximum visible in **Fig 1E**). Due to the curvature of the hippocampus and position of our electrode array, each shank sampled a slightly different depth of the CA1 lamina, but recordings could be localized based on physiological properties such as ripple frequency power, SWR CSD, and single unit locations (**Fig 1D**) to investigate layer-specific changes following TBI.

**Figure 1:**
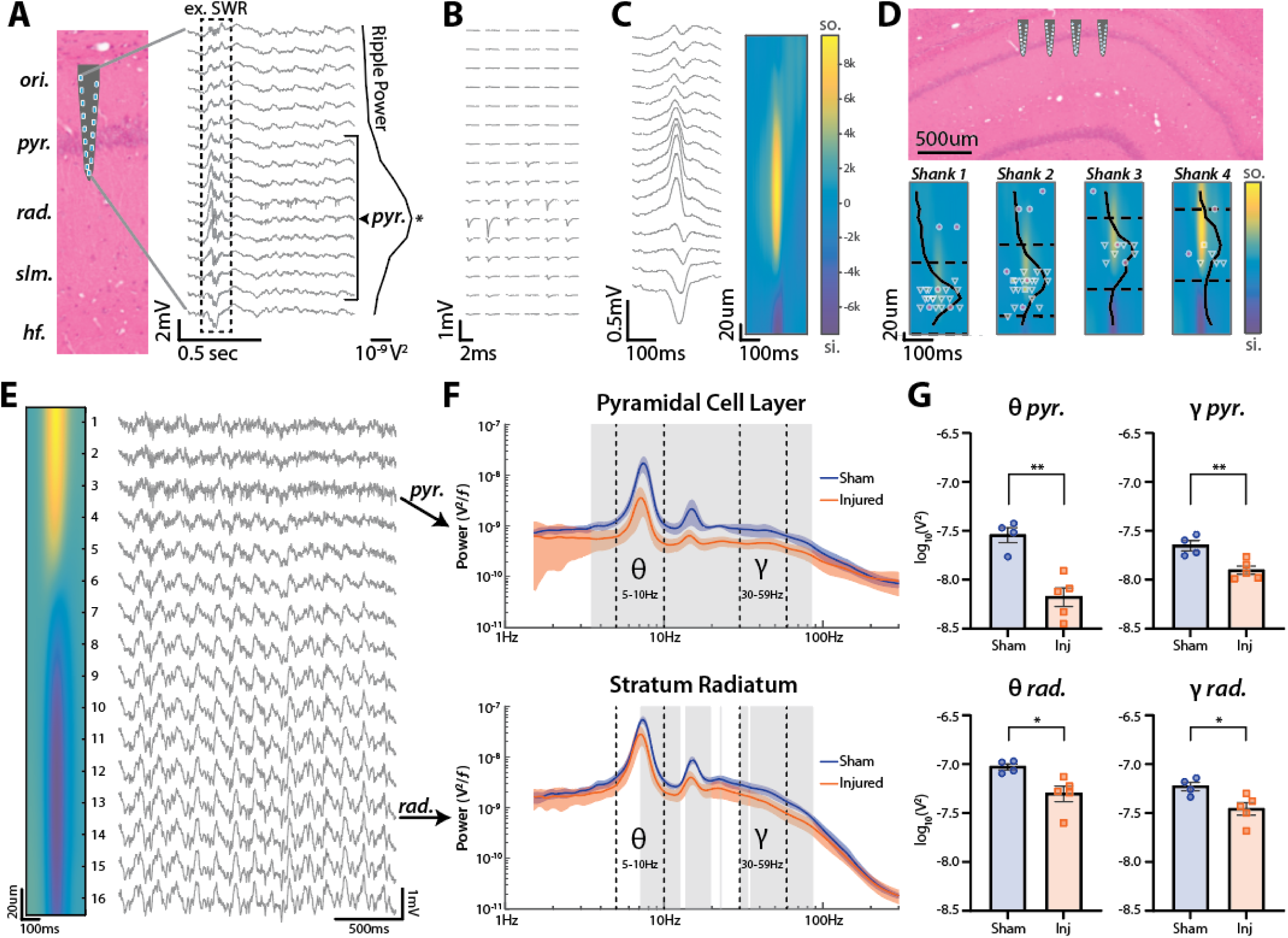
_L_FPI decreases theta and gamma power across CA1 lamina. **(A)** Silicon probe recording from dorsal CA1. Schematic of electrode shank widened for visualization but accurately scaled along CA1 lamina. The peak of ripple frequency (100-250 Hz) power across channels was used to define the *st. pyr* channel (marked with *). Channels within ±80 µm of the defined *st. pyr* channel were assigned to the pyramidal cell layer (denoted by bracket). **(B)** Mean waveforms from 6 single units identified as putative pyramidal cells recorded on the shank shown in A across all electrode contacts. **(C)** Mean SWR waveform from the shank shown in A and its associated current source density (CSD) **(D)** Schematic of an electrode array with shanks positioned along the proximal-distal axis of CA1. Heatmap contours show the mean SWR CSD for each shank. Normalized ripple frequency power and edges of the pyramidal cell layer are shown as solid and dashed lines respectively for each shank. Individual recorded units are shown at the depth of the channel containing their max amplitude (putative cell types: interneuron=magenta circle, pyramidal cell=green triangle, unclassified=yellow square). Shank 2 shown in A-C. **(E)** Example recording from a shank spanning both *st. pyr* and *st rad*. The peak of ripple frequency power occurs at channel 3 (identified as *pyr.*). The *st. rad* channel was defined as the channel closest to the peak sink in the CSD of the mean SWR waveform (channel 13 denoted by *rad.*). Note theta (5-10 Hz) and gamma (30-59 Hz) oscillations especially apparent in *st. rad*. **(F)** Power spectra (mean±SEM) from *st. pyr* (top) and *st. rad* (bottom) for sham and injured rats moving (>10 cm/sec) in the familiar environment. Theta (5-10 Hz) and gamma (30-59 Hz) bands denoted by dashed lines. Statistically significant differences between sham and injured are outlined in gray (α=0.01 for multiple comparisons; t-tests). **(G)** Integral of power in the theta and gamma frequency bands (mean±SEM; individual animals labeled by points) from sham and injured rats in *st. pyr* (top; theta: sham=-7.54±0.08 log_10_V^2^, injured=-8.18±0.09 log_10_V^2^, p=0.002; gamma: sham=-7.65±0.05 log_10_V^2^, injured=-7.90±0.04 log_10_V^2^, p=0.007; t-test) and *st. rad* (bottom; theta: sham=-7.03±0.03 log_10_V^2^, injured=-7.31±0.08 log_10_V^2^, p=0.022; gamma: sham=-7.23±0.04 log_10_V^2^, injured=-7.46±0.06 log_10_V^2^, p=0.028; t-test).

Hippocampal theta and gamma oscillations can be readily observed while rats actively explore an environment^51,56^, and these oscillations change in power and phase along the CA1 lamina (**Fig 1E**). Thus, precisely localizing electrodes is essential, especially when comparing across animals and groups. Because previous studies have reported either a broadband or frequency-specific reduction in oscillatory power following TBI^40,42,45^, we first took an agnostic approach and compared power spectra across sham and injured animals to determine whether there were layer-specific differences. Power spectra were computed from channels localized to *st. pyr* or *st. rad* while rats were actively moving (>10 cm/sec) in the familiar environment, averaged across shanks and recording days to obtain a mean for each animal, then averaged across sham and TBI conditions (**Fig 1F**). In *st. pyr*, injured rats had decreased power from 3.5-85.1 Hz, and in *st. rad*, they had decreased power in the 7.1-12.7, 13.7-19.7, 30.1-33.8, and 35.1-86.8 Hz frequency bands (t-tests; α=0.01 for multiple comparisons). Importantly, there was no significant difference in movement velocity between sham and injured animals (**Sup Fig 1**) indicating that reductions in power were not due to potential motor deficits in injured animals.

When separated into the well-characterized theta (5-10 Hz) and low-gamma (30-59 Hz) frequency bands, we found that TBI rats exhibited a significant reduction in both theta and low-gamma power in *st. pyr* (theta: p=0.002; gamma: p=0.007; t-test) as well as *st. rad* (theta: p=0.022; gamma: p=0.028 t-test; **Fig 1G**). These significant differences were also observed when animals were still (<10cm/sec; **Sup Fig 2**) and when a stricter value of >20 cm/sec was used as a threshold for moving (**Sup Fig 3**). Importantly, these theta and gamma frequency bands only comprise a portion of the broadband decrease in power observed in **Fig 1F**. Thus, we used a well-established method to flatten the power spectra^69^ in order to compare the theta and gamma frequency bands in the context of this broadband shift. Using this correction, we found that theta power was still reduced in injured animals in both *st. pyr* (p=0.002, t-test) and *st. rad* (p=0.021, t-test), but the reduction in gamma power was no longer statistically significant from either region (p>0.05, t-test; **Sup Fig 4**). It is important to note that power spectra from sham and injured rats converge at low (<4Hz) and high (>90Hz) frequencies. Thus, the decrease in power observed in injured rats is still limited in the frequency domain and does not represent a full-spectrum broadband shift. Taken together, our results demonstrate that injured rats have decreased oscillatory power spanning a range of physiological frequencies including theta and gamma, and that the loss of power is more pronounced in *st. pyr* than in *st. rad*, particularly in the theta frequency band.

**Figure 2:**
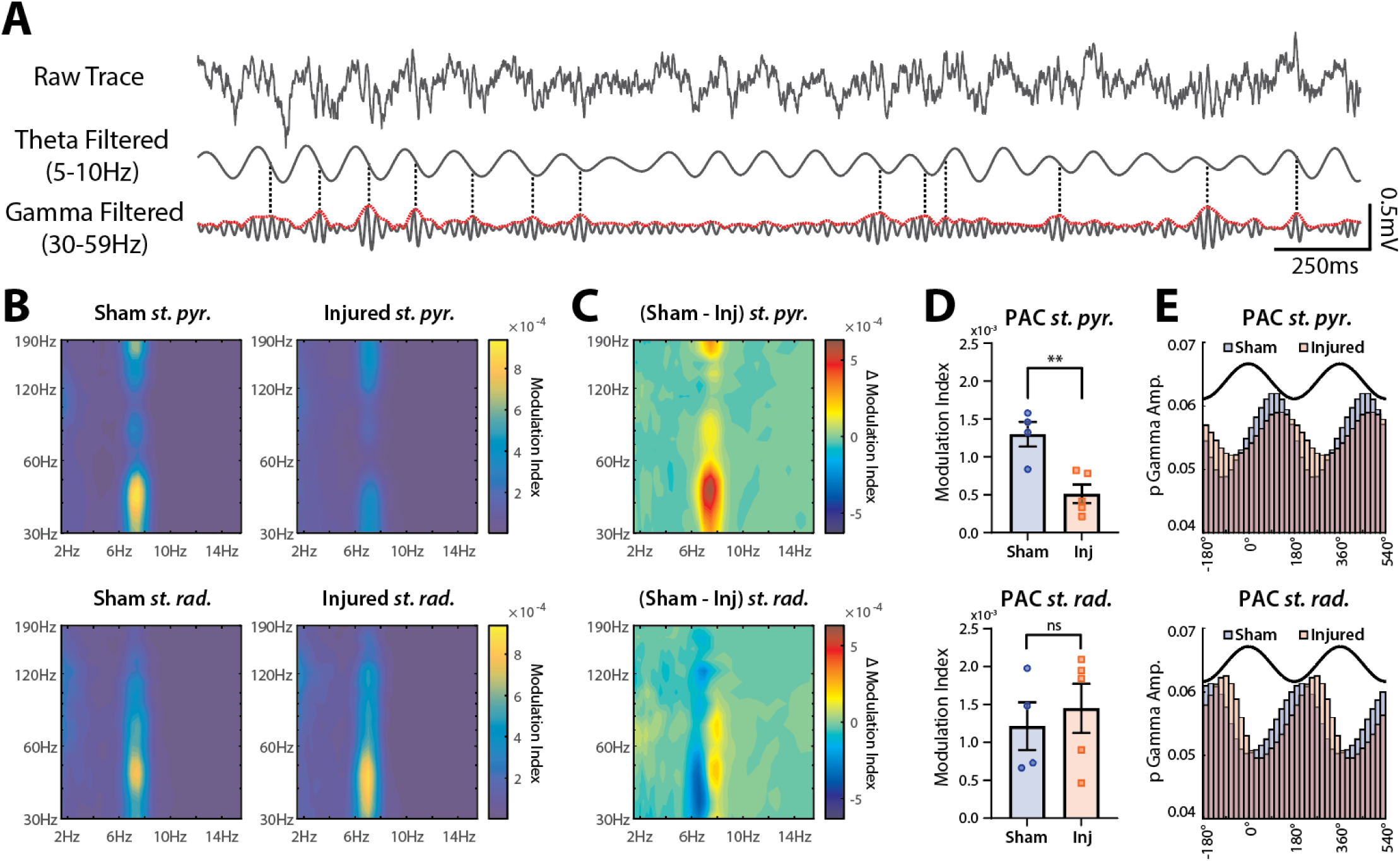
_L_FPI decreases theta-gamma phase-amplitude coupling (PAC) in st. pyr but not st. rad. **(A)** Visualization of theta-gamma PAC. Raw 3 sec trace recorded from *st. pyr* (top), the same trace filtered for theta (5-10 Hz; middle) and for gamma (30-59 Hz; bottom). Envelope amplitude of gamma is plotted in red and peaks in gamma amplitude >1SD above mean are marked with a dashed line. Note that most peaks align to a similar phase of theta (∼90°). **(B)** Averaged PAC heatmap contours from sham (left) and injured (right) animals in *st. pyr* (top) and *st. rad* (bottom). **(C)** Difference between sham and injured PAC heatmap contours in both *st. pyr* (top) and *st. rad* (bottom). **(D)** PAC modulation index values from broadband theta and gamma filtered signals (mean±SEM; individual animals labeled by points) in *st. pyr* (top; sham=0.00130±0.00016, injured=0.00051±0.00012, p=0.006, t-test) and *st. rad* (bottom; sham=0.00121±0.00032, injured=0.00145±0.00032, p=0.623, t-test). **(E)** Normalized gamma amplitudes across theta phase bins (2 cycles of theta shown in black for reference). Peak gamma in *st. pyr* (sham=89.2°, injured=115.2°, Δ=26°) and *st. rad* (sham=200.0°, injured=255.4°, Δ=55.4°).

**Figure 3:**
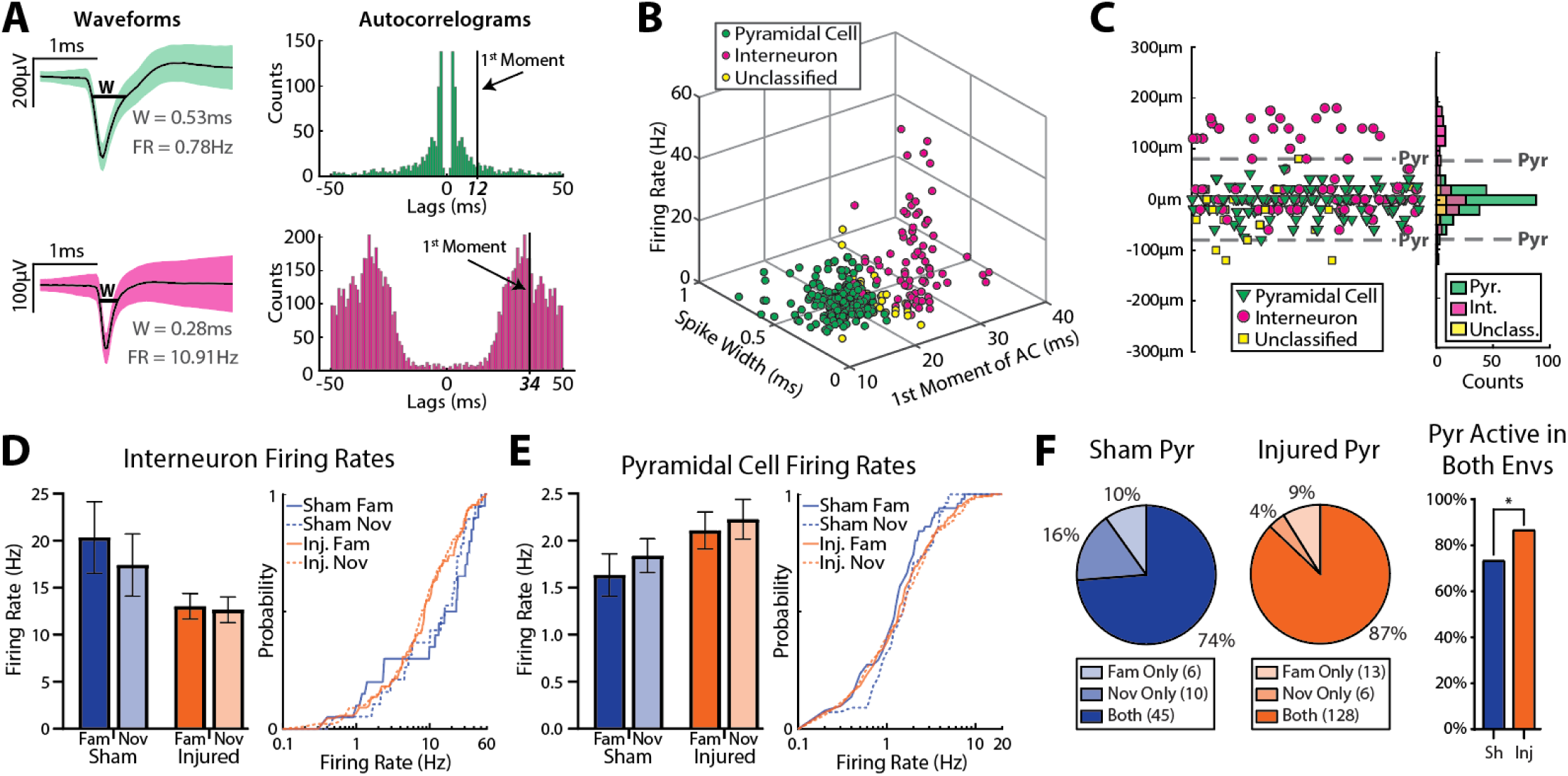
_L_FPI minimally affects single unit firing rates but increases recruitment of pyramidal cells. **(A)** Waveforms (mean±SD) from an example putative pyramidal cell (top) and interneuron (bottom) and their associated autocorrelograms (peak at 0ms omitted for clarity). W=width, FR=firing rate **(B)** Scatterplot of spike width, first moment of the autocorrelogram, and firing rate of all cells located within the pyramidal cell layer from both sham and injured animals. Note separation of putative pyramidal cells and interneurons with unclassified cells typically lying between these 2 populations. **(C)** Location of all cells relative to the defined *st. pyr* channel (based on peak ripple frequency power). Note the peak of the pyramidal cell distribution is located on the defined *st. pyr* channel, and the bimodal distribution of interneurons reflects populations localized to *st. oriens* and *st. pyr*. **(D)** Left: firing rates (mean±SEM) of interneurons in the familiar and novel environments across sham and injured animals (familiar: sham=20.3±3.8 Hz, n=20, injured=13.0±1.3 Hz, n=88, p=0.116; novel: sham=17.4±3.3 Hz, n=19, injured=12.6±1.4 Hz, n=89, p=0.185; ks-test). Right: cumulative distributions of interneuron firing rates. **(E)** Left: firing rates (mean±SEM) of pyramidal cells in the familiar and novel environments across sham and injured animals (familiar: sham=1.63±0.23 Hz, n=51, injured=2.11±0.20 Hz, n=141, p=0.446; novel sham=1.84±0.18 Hz, n=55, injured=2.23±0.21 Hz, n=134, p=0.170, ks-test). Right: cumulative distributions of pyramidal cell firing rates. **(F)** Recruitment of pyramidal cells across environments (sham=74%, injured=87%, p=0.025, Fisher’s exact test).

**Figure 4:**
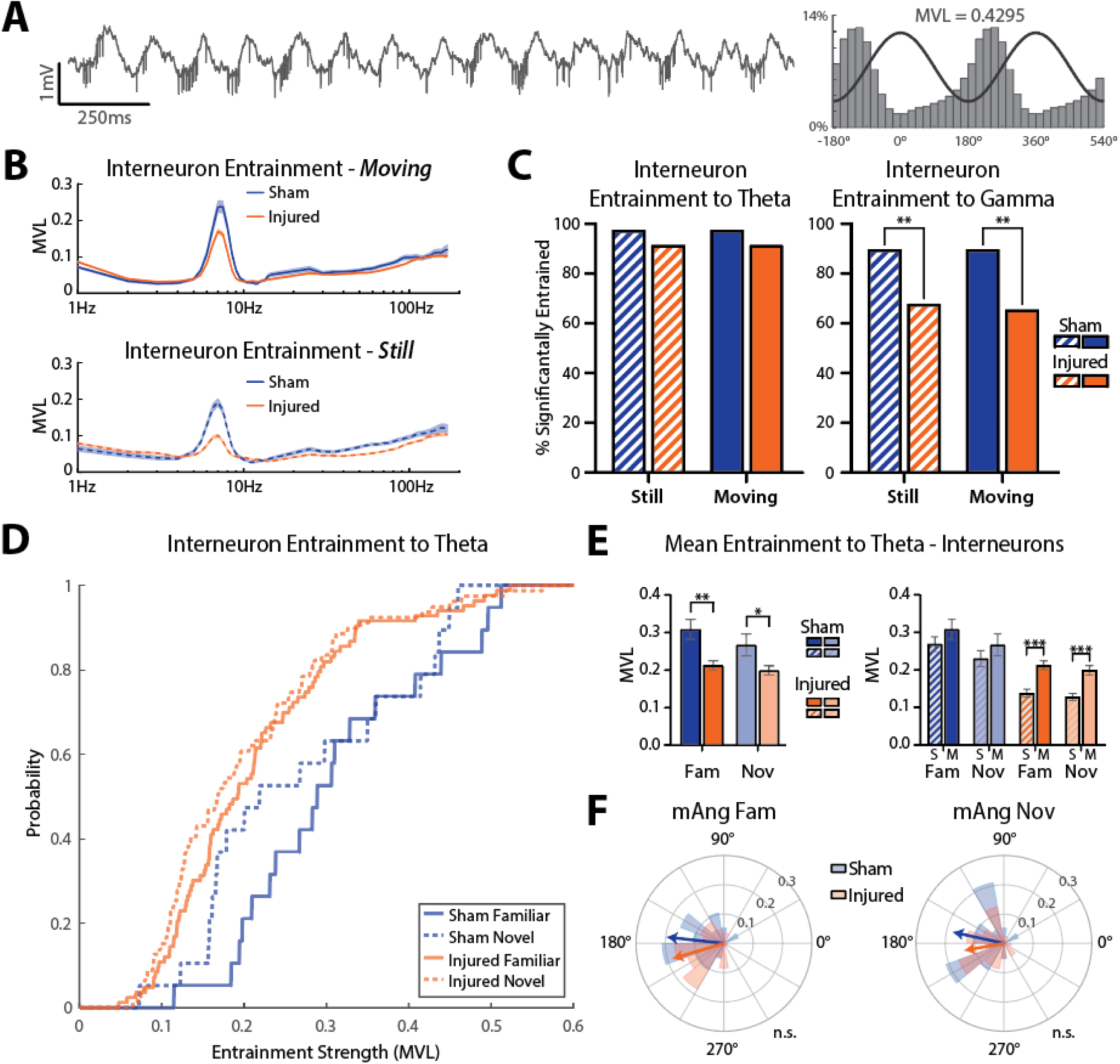
_L_FPI disrupts spike field coherence to hippocampal oscillations. **(A)** Example 2 sec recording trace containing a large amplitude interneuron entrained to theta (all spikes are from the same unit). Right: probability of spike times across theta phase bins (line shows 2 cycles of theta for reference). The mean vector length of this cell is 0.43. **(B)** Entrainment strength of all interneurons across frequencies in both environments while rats were moving (>10 cm/sec; top) or still (<10 cm/sec; bottom). **(C)** Percent of interneurons significantly entrained to theta (5-10 Hz) and gamma (30-59 Hz) while animals were still or moving (theta still: sham=97.44%, injured=91.53%, p=0.315; theta moving: sham=97.44%, injured=91.53%, p=0.315; gamma still: sham=89.74%, injured=67.80%, p=0.006; gamma moving: sham=89.74%, injured=65.54%, p=0.002; Fisher’s exact test). **(D)** Mean vector length cumulative distributions of all interneurons significantly entrained to theta across all conditions. **(E)** Left: average entrainment strength (mean±SEM) of interneurons significantly entrained to theta while rats were moving in the familiar (sham=0.31±0.03, n=19, injured=0.21±0.01, n=83, p=0.004, ks-test) and novel (sham=0.27±0.03, n=19, injured=0.20±0.01, n=79, p=0.026, ks-test) environment. Right: average entrainment of significantly entrained interneurons while rats were still or moving in each environment (sham familiar: still=0.27±0.02, n=19, moving=0.31±0.03, n=19, p=0.462; sham novel: still=0.23±0.02, n=19, moving=0.27±0.03, n=19, p=0.462; injured familiar: still=0.14±0.01, n=79, moving=0.21±0.01, n=83, p<0.001; injured novel: still=0.13±0.01, n=83, moving=0.20±0.01, n=79, p<0.001; ks-test). **(F)** Polar histograms showing the mean angle of entrainment for all significantly entrained interneurons while animals were moving in the familiar or novel environment (familiar: sham=173.7°, injured=196.6°, p=0.446; novel: sham=167.6°, injured=190.2°, p=0.443; circular Kruskal-Wallis test). All theta is referenced to the defined *st. pyr* channel on the shank from which units were recorded (peak of theta is 0°).

### _L_FPI decreases theta-gamma PAC in st. pyr but not st. rad

Theta-gamma PAC (**Fig 2A**) is a well-characterized phenomenon important for the selection and coordination of cell ensembles that support spatial coding and memory processes^56,59,60^. PAC requires temporal coupling of afferent input with local inhibition and is differentially observed across the layers of CA1^47,59^. Importantly, while the generation and coupling of oscillations are likely to involve partially overlapping mechanisms, a loss of oscillatory power does not necessarily result in reduced PAC. Because we expect TBI to induce disruptions in network synchrony^70^, we examined whether theta-gamma PAC would be reduced following TBI. To assess the effects of TBI on theta-gamma PAC, we computed the modulation index^71^ across discrete frequency bands during epochs in which rats were moving (>10cm/sec) in the familiar environment, then constructed PAC heatmaps for channels localized to *st. pyr* and *st rad* (**Fig 2B**). Sham rats showed strong theta-gamma PAC in the pyramidal cell layer which was drastically reduced in injured rats. This reduction was specific to *st. pyr*, as the strength of PAC in *st. rad* was similar between sham and injured rats, though there was a slight shift in coupling to a lower frequency of phase and amplitude. Both the loss of PAC strength in *st. pyr* and the frequency shifts in *st. rad* can be visualized by subtracting PAC heatmaps of injured animals from sham animals (**Fig 2C**).

To quantify differences in theta-gamma PAC between sham and injured rats, we extracted the phase and amplitude from broadband theta (5-10 Hz) and low-gamma (30-59 Hz) filtered signals respectively and compared the magnitude and phase preference of PAC. We found that the magnitude of theta-gamma PAC in injured rats was reduced in *st. pyr* (p=0.006, t-test) but not in *st. rad* (p=0.623, t-test; **Fig 2D**). Additionally, the maximal amplitude of gamma was coupled to a later phase of theta in injured rats compared to shams in both *st. pyr* (sham=89.2°, injured=115.2°, Δ=26.0°; theta peak=0°) and *st. rad* (sham=200.0°, injured=255.4°, Δ=55.4°; **Fig 2E**). Overall, injured rats show a decrease in the strength of theta-gamma PAC that is specific to *st. pyr*, and a shift in the peak gamma amplitude to a later phase of theta in both *st. pyr* and *st. rad*, which suggests a potential desynchronization of long-range theta inputs with local gamma processing in CA1.

### _L_FPI minimally affects single unit firing rates but increases recruitment of pyramidal cells

Oscillations in CA1 largely reflect synchronous synaptic inputs to this region, which are known to drive cells to fire in a temporally precise manner. The reduction in oscillatory power and coupling following TBI suggests that single unit recruitment and spike timing during behavior may also be disrupted. Thus, we compared neuronal firing properties between sham and injured animals by isolating single units and clustering them into one of 3 categories: putative pyramidal cells, putative interneurons, and unclassified (see Methods; putative identifiers henceforth removed for writing clarity). Because both pyramidal cells and interneurons are present in the pyramidal cell layer, units within this layer (±80 µm of the defined *st. pyr* channel) were manually clustered based on each unit’s spike width, firing rate, and 1^st^ moment of the autocorrelogram (a measure of burstiness) as previously described^72^ (**Fig 3A,B**). Pyramidal cells typically have wider waveforms, lower firing rates and are more bursty (lower 1^st^ moment of the autocorrelogram) compared to interneurons. Thus, these two populations form distinct clusters when these features are plotted together (**Fig 3B**). Manual clustering was compared to automated k-means consensus clustering and there was a 95.47% overlap between clustering methods when the unclassified cell group was omitted (**Sup Fig 5**). We chose to use manual clustering because the inclusion of the unclassified group allowed for a more conservative classification, and manual clustering is more robust to outliers than automated clustering (example in **Sup Fig 5**). Cells located >80 µm above the defined *st. pyr* channel were all classified as interneurons as anatomically these cells are located in *st. oriens* where there are no pyramidal cells. Importantly, cells classified as interneurons based on their anatomical location had firing properties that clustered with other interneurons further supporting their classification (**Sup Fig 6**). When the location of each shank was centered to the defined *st. pyr* channel, cells are visibly distributed into the distinct *st. oriens* and *st. pyr* cell layers (**Fig 3C**).

**Figure 5:**
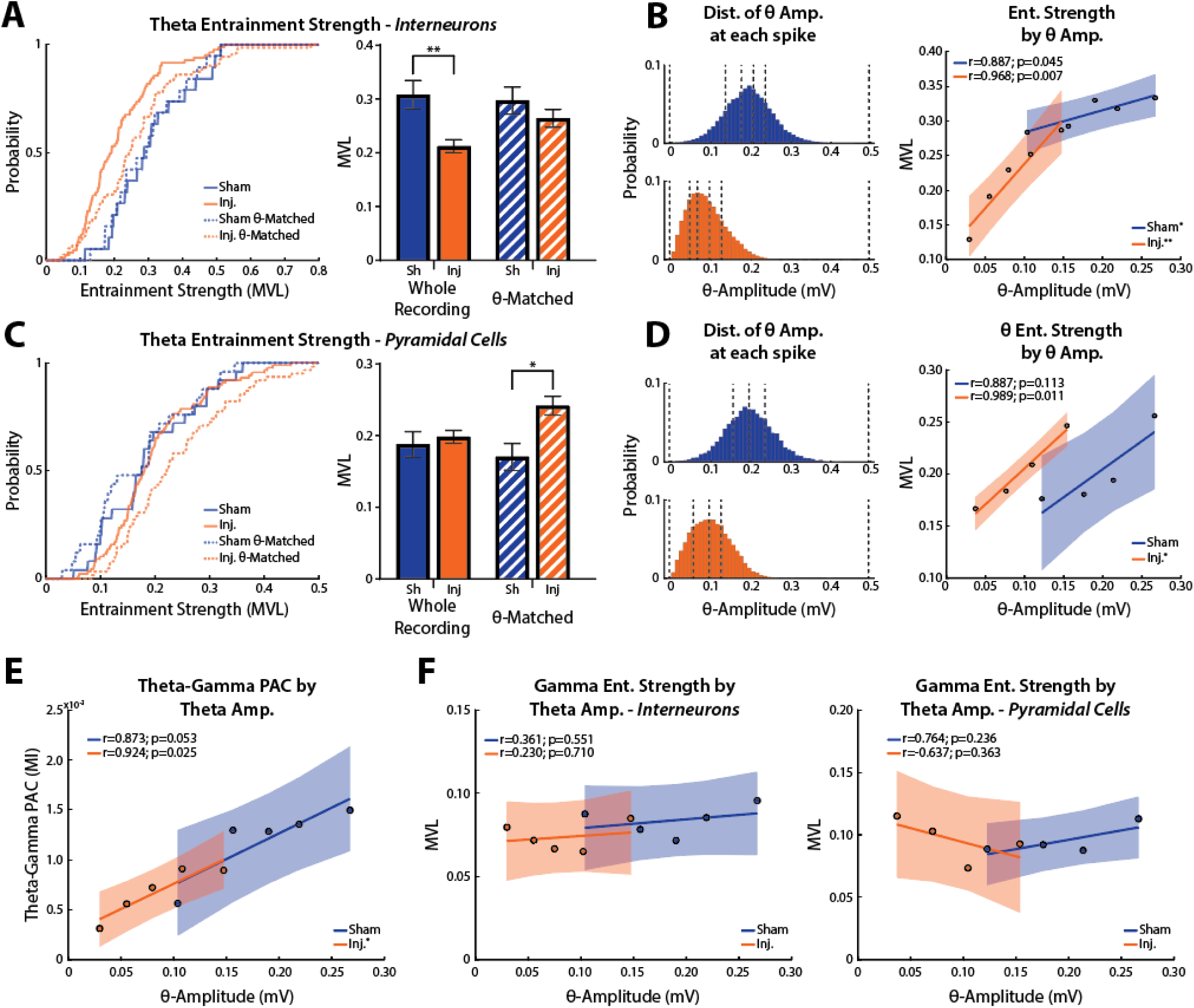
Theta amplitude is coupled to single unit entrainment to theta and PAC in injured animals. **(A)** Left, cumulative distributions of interneuron theta entrainment strength (MVL) for the entire recording (solid lines) and during periods of matched theta power (dashed lines). Right, average (mean±SEM) entrainment strength for each condition (entire recording: sham=0.31±0.03, n=19, injured=0.21±0.01, n=83, p=0.004; theta-matched: sham=0.30±0.03, n=19, injured=0.26±0.02, n=72, p=0.313; ks-test). Significantly entrained cells only. **(B)** Left, distribution of theta amplitudes at every interneuron spike from sham (top) and injured (bottom) animals. Distributions are split into 5 bins with an equal number of spikes in each bin (dashed lines denote bin edges). Right, theta entrainment strength averaged across neurons for each theta amplitude bin (linear regression fit line with shaded 95% confidence interval). The entrainment strength of interneurons from both sham and injured animals is correlated with theta amplitude (sham: r=0.887, slope=0.32, p=0.045; injured: r=0.968, slope=1.28, p=0.007). **(C)** Left, cumulative distributions of pyramidal cell theta entrainment strength for the entire recording (solid lines) and during periods of matched theta power (dashed lines). Right, average (mean±SEM) theta entrainment strength for each condition (entire recording: sham=0.19±0.02, n=25, injured=0.20±0.01, n=89, p=0.518; theta-matched: sham=0.17±0.02, n=25, injured=0.24±0.01, n=62, p=0.012; ks-test). Significantly entrained cells only. **(D)** Left, distribution of theta amplitudes at every pyramidal cell spike from sham (top) and injured (bottom) animals. Distributions are split into 4 bins with an equal number of spikes in each bin (dashed lines denote bin edges). Right, theta entrainment strength averaged across neurons for each theta amplitude bin (linear regression fit line with shaded 95% confidence interval). The entrainment strength of pyramidal cells from injured but not sham animals is correlated with theta amplitude (sham: r=0.887, slope=0.54, p=0.113; injured: r=0.989, slope=0.69, p=0.011). **(E)** Theta-gamma PAC binned using the same theta amplitude bins as in B. There was a correlation between theta amplitude and theta-gamma PAC in injured (r=0.924, slope=0.005, p=0.025) but not sham animals (r=0.873, slope=0.005, p=0.053). **(F)** Strength of entrainment to gamma in interneurons (left) and pyramidal cells (right) at the same theta amplitude bins as B and D. In both sham and injured animals, there was no correlation between theta amplitude and gamma entrainment strength in interneurons (sham: r=0.361, slope=0.05, p=0.551; injured: r=0.230, slope=0.04, p=0.710) or in pyramidal cells (sham: r=0.764, slope=0.15, p=0.236; injured: r=-0.637, slope=-0.23, p=0.363). All data presented is while animals were moving in the familiar environment.

**Figure 6:**
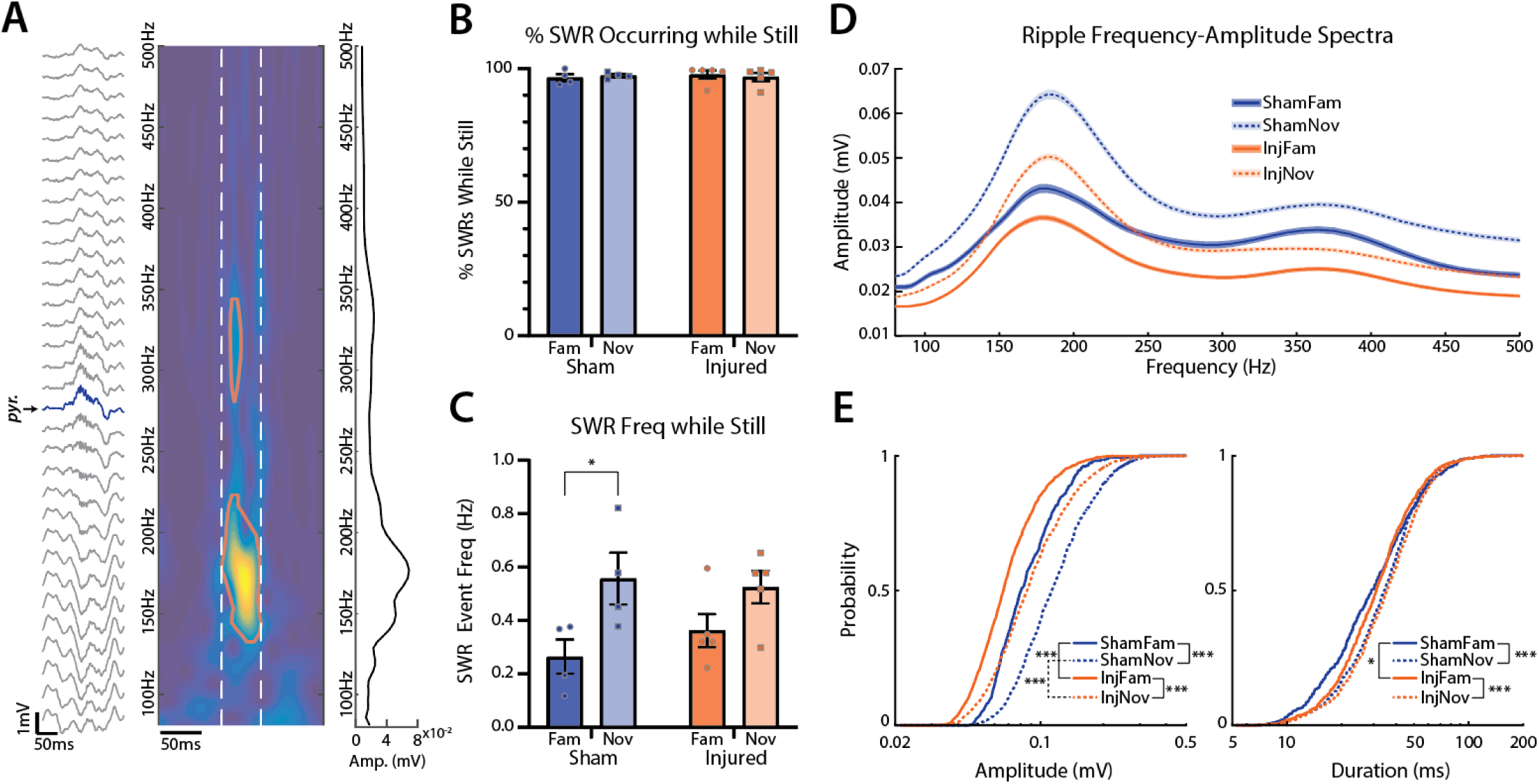
_L_FPI decreases SWR amplitude. **(A)** Example SWR event recorded across all electrode contacts from a single shank (*st. pyr* channel labeled and colored blue), the associated time-frequency map from the *st. pyr* channel (red contour outlines where ripple event exceeds amplitude threshold; white dashed lines mark edges of ripple event in time), and the frequency-amplitude spectrum of the event averaged over time between white dashed lines. **(B)** Percentage of SWRs that were detected while the animal was still (sham: familiar=96.7±0.7%, novel=97.5±0.3%, injured: familiar=97.9±0.7%, novel=96.9±0.7%; mean±SEM; points represent individual animals). **(C)** SWR event rate normalized to the amount of time animals were still (sham: familiar=0.26±0.03 Hz, novel=0.56±0.05 Hz, p=0.011; injured: familiar=0.36±0.03 Hz, novel=0.52±0.03 Hz, p=0.052; t-test; mean±SEM; points represent individual animals). **(D)** Average (mean±SEM) frequency-amplitude spectrum of all ripples across conditions. **(E)** Left: cumulative distributions of ripple amplitudes across conditions (familiar: sham=91.2±1.7 mV, n=490, injured=73.3±0.9 mV, n=1082, p<0.001; novel: sham=126.2±1.6 mV, n=1285, injured=98.4±1.3 mV, n=1464, p<0.001; sham still vs moving: p<0.001; injured still vs moving: p<0.001; mean±SEM; ks-test). Right: cumulative distributions of ripple durations across conditions (familiar: sham=34.3±0.9 ms, injured=34.7±0.6 ms, p=0.019; novel: sham=36.9±0.5 ms, injured=38.5±0.5 ms, p=0.062; sham still vs moving: p<0.001; injured still vs moving: p<0.001; mean±SEM; ks-test).

Hyperexcitability in CA1 is a proposed mechanism for hippocampal dysfunction, but there are contradicting reports about the effects of TBI on the firing rates of individual neurons^5,41,43,44,73^. We therefore assessed whether TBI broadly affects firing properties or overall recruitment of these specific cell populations across different conditions. Specifically, we evaluated whether the changes in oscillatory power and coupling coincided with changes in single unit firing rates or overall recruitment across distinct environments. We compared the firing rates of active (firing rate >0.1 Hz) pyramidal cells and interneurons in the familiar and novel environment and found no significant differences in the firing rate of interneurons (familiar: p=0.116; novel: p=0.185; ks-test; **Fig 3D**) or pyramidal cells (familiar: p=0.446; novel: p=0.170; ks-test; **Fig 3E**) between sham and injured groups. Overall, firing rates were significantly higher when rats were moving (>10 cm/sec) compared to when they were still, but there were no significant differences in firing rates between the sham and injured groups in either the still or moving condition (**Sup Fig 7**). Additionally, there were no differences in pyramidal cell bursting between sham and injured animals in either environment or across still and moving conditions (**Sup Fig 8**), which suggests that intrinsic excitability has not changed, or does not drive changes in firing rates between groups. Interestingly, a higher percentage of pyramidal cells were active (firing rate >0.1Hz) in both the familiar and novel environment in injured rats compared to shams (sham=74%, injured=87%, p=0.025, Fisher’s exact test; **Fig 3F**). This suggests that rather than affecting the baseline firing properties of individual cells, which may remain relatively stable due to homeostatic plasticity mechanisms, TBI-associated changes in CA1 circuit excitability can result in an over-recruitment of pyramidal cells across distinct conditions.

### _L_FPI disrupts spike field coherence and impairs single unit entrainment to theta oscillations

While the overall firing rates of single units were not significantly altered in TBI rats, the observed reductions in oscillatory power and PAC implicate disrupted synchronization of single unit firing. Many CA1 neurons are entrained to local oscillations and are more likely to fire around a specific phase of the oscillation (example of an interneuron entrained to theta shown in **Fig 4A**). Proper entrainment of interneurons is especially important for temporal coding as they support the selection and temporal coordination of behaviorally relevant cell ensembles. In order to determine whether TBI disrupts single unit entrainment to local oscillations, we first assessed entrainment strength by calculating the mean vector length (MVL) of interneuron spike times across a range of discrete frequency bands (extracted from the defined *st. pyr* channel on the electrode shank from which neurons were recorded) while rats were either still (<10 cm/sec) or moving (>10 cm/sec; **Fig 4B**). Clear peaks in entrainment can be seen in the theta frequency range for both conditions.

To quantify single unit entrainment to physiologically relevant hippocampal oscillations, we calculated the percentage of interneurons that were significantly entrained to broadband filtered theta (5-10 Hz) and gamma (30-59 Hz) oscillations recorded from the *st. pyr* channel using circular shuffling (see methods). There was no difference in the percentage of interneurons significantly entrained to theta between sham and injured rats when animals were still (sham=97.44%, injured=91.53%, p=0.315, Fisher’s exact test) or moving (sham=97.44%, injured=91.53%, p=0.315, Fisher’s exact test). However, a lower percentage of interneurons were entrained to gamma in injured rats compared to shams both while animals were still (sham=89.74%, injured=67.80%, p=0.006, Fisher’s exact test) and moving (sham=89.74%, injured=65.54%, p=0.002, Fisher’s exact test; **Fig 4C**). While a similar percentage of interneurons were significantly entrained to theta in both sham and injured animals, the reduced peak in **Fig 4B** suggests that interneurons from injured rats may be less strongly entrained to theta. To investigate this possibility, we extracted only interneurons that were significantly entrained to theta and compared their entrainment strength and phase preference in both environments. **Figure 4D** shows entrainment strength cumulative distributions for interneurons significantly entrained to theta while rats were moving. Interneurons were less entrained to theta in injured rats compared to shams in both the familiar (p=0.004, ks-test) and novel (p=0.026, ks-test) environment (**Fig 4E**). Additionally, the MVL of interneurons was dramatically higher when injured rats were moving compared to when they were still (familiar: p<0.001; novel: p<0.001; ks-test), which was not the case in sham rats (familiar: p=0.462; novel: p=0.462; ks-test; **Fig 4E**). Despite the reduction in entrainment strength, the mean angle of entrainment for interneurons significantly entrained to theta was not different between sham and injured rats in either environment while animals were moving (familiar: sham=173.7°, injured=196.6°, p=0.446; novel: sham=167.6°, injured=190.2°, p=0.443; circular Kruskal-Wallis test; theta peak=0°; **Fig 4F**).

Overall, while interneurons were entrained to the same angle of theta in both groups, interneurons from injured animals fired more promiscuously around this angle and were also less entrained to gamma. Because interneurons are known to strongly influence spike timing in pyramidal cells, these results reveal potential disruptions to the generation and temporal coordination of oscillations that support information processing in the hippocampus.

### Theta power is coupled to entrainment strength and PAC in injured animals

Disentangling the generation of oscillations and single unit entrainment to oscillations in the hippocampus is complicated due to emergent network dynamics in the hippocampus and the multiple contributors to oscillatory power. Because these processes are thought to be interdependent, it is difficult to mechanistically untangle the loss of theta power and reduced entrainment strength of interneurons to theta in injured animals. On one hand, the reduced entrainment strength of CA1 interneurons could directly drive the observed decrease in theta power, implicating local interneurons and/or their upstream inputs in TBI-associated theta loss. Alternatively, the loss of theta power could occur independently from the decreased entrainment strength of interneurons, which would implicate TBI-associated disruptions to alternative theta generators. In order to help distinguish between these possibilities, we subsampled the data into periods of time in which theta power was comparable between sham and injured animals and compared entrainment strength specifically at these times (see Methods). We found that during periods of comparable theta power, interneuron entrainment strength was similar between sham and injured animals (p=0.313, ks-test; **Fig 5A**). To further investigate the relationship between theta power and interneuron entrainment strength, we created a distribution of theta amplitudes at each interneuron spike for sham and injured animals, divided these distributions into equally sized bins, and computed the entrainment strength in each bin (**Fig 5B**; see Methods). We found a weak but statistically significant correlation between theta amplitude and interneuron entrainment strength in sham animals (r=0.887, p=0.045) that was much more pronounced in injured rats (r=0.968, p=0.007; **Fig 5B**). Together, these results suggest a direct link between the reduction in theta power and interneuron entrainment strength in injured animals, potentially implicating CA1 interneurons or their upstream inputs in network dysfunction.

In addition to contributing to theta generation, interneurons are known to constrain the active population of CA1 pyramidal cells and strongly control their spike timing^74–76^. Pyramidal cells fire more flexibly across different phases of theta compared to interneurons, and place cells are known to exhibit phase precession in which they fire at earlier phases of theta as animals pass through their place field. This is thought to contribute to spatial coding, as phase precession is important for generating theta sequences that can be predictive of behavior^37,77–81^. Thus, the flexible firing of pyramidal cells across theta phases is proposed to be important for spatial learning and memory. Because interneurons tightly control pyramidal cell firing and exhibit changes in entrainment strength in TBI rats, we assessed whether pyramidal cells fired flexibly across phases of theta, or whether they were more entrained to specific phases following TBI. Additionally, due to the tight relationship between theta power and interneuron entrainment strength in TBI animals, we also assessed whether there were differences in entrainment strength during epochs of matched theta power. At baseline, we found no difference in the entrainment strength of pyramidal cells between sham and injured rats (p=0.518, ks-test; **Fig 5C**). However, we found that pyramidal cells from injured rats were more strongly entrained to theta than in sham rats during periods of matched theta power (p=0.12, ks-test; **Fig 5C**). Additionally, we found a strong correlation between theta amplitude and the entrainment strength of pyramidal cells in injured but not sham animals when data was subsampled as described for interneurons (sham: r=0.887, p=0.113; injured: r=0.989, p=0.011; **Fig 5D**). This demonstrates that while pyramidal cells from sham and injured animals fire flexibly across the theta cycle, periods of increased theta power are associated with less flexible firing of pyramidal cells in injured animals which may impair the generation of theta sequences that support behavior.

Flexible firing of pyramidal cells across the theta cycle is also important because memory encoding and retrieval are thought to be biased towards different phases of theta when LTP is strongest (peak) or weakest (trough) respectively, a process supported by theta-gamma PAC^48^. Because both pyramidal cells and interneurons from injured animals showed a strong relationship between theta amplitude and entrainment, we assessed whether the strength of theta-gamma PAC was also correlated with theta amplitude in injured animals. Using the same theta amplitude bins used for assessing interneuron entrainment strength (**Fig 5B**), we found that the strength of PAC was correlated with theta amplitude in the injured (r=0.924, p=0.025) but not sham (r=0.873, p=0.053) animals (**Fig 5E**). Because gamma oscillations are generated locally through a mix of interneuron and pyramidal cell-interneuron network interactions, we assessed whether single unit entrainment to gamma was also positively correlated with theta amplitude in injured animals as it was for PAC. Surprisingly, gamma entrainment strength of both pyramidal cells and interneurons was not correlated with theta amplitude in either sham (interneurons: r=0.361, p=0.551; pyramidal cells: r=0.764, p=0.236) or injured (interneurons: r=0.230, p=0.710; pyramidal cells: r=-0.637, p=0.363) animals (**Fig 5F**).

Overall, interneurons from injured animals were less entrained to theta compared to shams, but pyramidal cells from both groups fired flexible across the theta cycle. However, with increasing theta amplitude, both interneurons and pyramidal cells from injured animals exhibited increased entrainment to theta, and at comparable theta amplitudes, there was no difference in the entrainment strength of interneurons between groups, but pyramidal cells from injured animals were more strongly entrained to theta than in shams. Theta-gamma PAC also increased with theta amplitude in injured animals but single unit entrainment to gamma did not.

### _L_FPI decreases SWR amplitude

CA1 SWRs are highly coordinated events implicated in memory consolidation and planning behaviors^65^. These events require precise synchronization of distributed cell populations resulting in a sharp-wave sink in *st. rad* with an associated ripple oscillation (150-250 Hz) in *st. pyr*^64^. The TBI-associated decrease in power, PAC, and entrainment that occurred while animals were actively moving suggests an overall desynchronization of hippocampal activity in the active state, thus we predicted that TBI would also disrupt highly coordinated SWRs in CA1 during quiet wakefulness. To assess this, putative SWRs were automatically detected by identifying candidate ripple events (using time-frequency decomposition as described previously^82^) that coincided with a sharp wave, then manually curated by a blinded observer (see Methods). **Figure 6A** shows an example SWR event along with its time-frequency map. We found that the vast majority of detected SWR events occurred while animals were still (sham: familiar=96.7±0.7%, novel=97.5±0.3%; injured: familiar=97.9±0.7%, novel=96.9±0.7%; mean±SEM; **Fig 6B**), which is expected, as SWRs are known to preferentially occur during awake immobility^64,65,82^.

To evaluate whether TBI disrupts ripple generation, we assessed SWR event frequency between sham and injured rats across environments normalized to the total amount of time that rats were still (**Fig 6C**). While there were no significant differences between sham and injured animals, the rate of SWR events was significantly higher in the novel compared to the familiar environment in sham (p=0.011, t-test) but not injured (p=0.052, t-test) animals, likely due to the slightly higher rate of SWR events in injured animals in the familiar environment. **Figure 6D** shows the spectral profile of ripples detected in *st. pyr* across conditions. As expected, ripple amplitudes and durations (**Fig 6E**) were higher in the novel compared to familiar environment in sham animals (amplitude: p<0.001; duration: p<0.001; ks-test), and we found the same was true for injured animals (amplitude: p<0.001; duration: p<0.001; ks-test). Ripple durations were slightly longer in injured rats compared to shams, but this difference only reached statistical significance in the familiar (p=0.019, ks-test) but not novel (p=0.062) environment (**Fig 6E**). However, ripple amplitudes were greatly reduced in injured animals compared to shams in both environments (familiar: p<0.001; novel: p<0.001; ks-test; **Fig 6E**). Together, these results demonstrate that a dynamic range of ripple amplitudes and durations still exists across environmental conditions in injured animals, but the TBI-associated decrease in ripple power suggests that TBI disrupts ripple generation mechanisms, which are known to support memory consolidation.

### _L_FPI disrupts spatial memory and induces stereotypic pathologies

The _L_FPI model of TBI in rats is known to produce spatial memory deficits and stereotypic patterns of pathology^83–87^, however, _L_FPIs can vary significantly across studies. Injury severity can be titrated based on the intensity of the fluid pulse generated, but other factors such as the precise placement of the craniectomy as well as differences in surgical technique and injury devices used can all contribute to _L_FPI heterogeneity^88,89^. Because _L_FPIs are not always standardized, we assessed whether _L_FPIs from this study produced pathology in line with previously published results from this model and reproduced spatial memory deficits^43,86,90^.

Spatial memory following TBI was assessed in a separate cohort of rats using the Morris Water Maze (MWM) as described previously^86^ (see methods). Animals were trained to find a submerged hidden platform in a pool then were subjected to a _L_FPI (1.34-1.80 atm) or sham surgery. Two days later, rats were placed back into the pool to test their memory of the platform location. On test day, injured rats had a lower memory score than shams (p=0.020, Welch’s t-test) indicating poor spatial memory (**Sup Fig 9A**). Importantly, the swim speed and distance traveled were not different between sham and injured rats during testing (**Sup Fig 10A**) indicating that differences in memory scores were not due to potential motor deficits from TBI. Additionally, the learning curves for each group were the same before sham/injury surgery (**Sup Fig 10B**).

A subset of 1 sham and 2 injured rats were sacrificed immediately after testing in the MWM (at 48hrs post _L_FPI/sham surgery) to verify that pathology generated in this study was in line with previously published results. Injured animals demonstrated multifocal hemorrhagic lesions including within the corpus callosum directly underlying the impact site, the angular bundle, fimbria-fornix and the lateral temporoparietal cortex (**Sup Fig 9B**). In one animal the temporoparietal hemorrhage extended into the lateral aspect of the ipsilateral hippocampus. APP immunoreactive axonal pathology was also observed consistently post-injury. Injured axons displayed abnormal morphologies including varicose swellings and axon bulbs indicative of transport interruption^91–94^ (**Sup Fig 9C,D**). Consistent with prior descriptions of the model^43^, axonal pathology was observed in a multifocal distribution post-injury including within the perilesional cortex, corpus callosum, thalamus, angular bundle (**Sup Fig 9C**), and fimbria-fornix (**Sup Fig 9D**). Notably, the fimbria-fornix and angular bundle serve as two major input/output pathways connecting the hippocampus to other structures. Degenerating cells, identified via Fluoro-Jade C staining were observed within the peri-lesional cortex and thalamus (**Sup Fig 9F**). While cell degeneration was also identified in the dentate granule cell layer in both animals, no cell degeneration was observed in CA1 (**Sup Fig 9G**) or hilus of the hippocampus, or entorhinal cortex. In one injured animal, minimal neuronal cell death was observed in CA2-3. No axonal pathology, cell death or hemorrhage was observed in the sham animal in any region examined (**Sup Fig 9E,H**).

## DISCUSSION

Learning and memory deficits are common following TBI, however it remains poorly understood how TBI-associated pathological changes disrupt information processing in brain regions important for these cognitive processes. Therefore, we utilized high-density electrophysiology to record from the hippocampus in freely moving rats following the well-characterized _L_FPI model of TBI. We took advantage of laminar probes to characterize layer-specific pathophysiology across CA1 lamina following TBI and found a loss of theta and gamma power across *st. pyr* and *st. rad* in injured animals, along with a drastic reduction in PAC that was specific to *st. pyr*. Interneurons from TBI rats were less entrained to theta and gamma, but their theta entrainment strength increased as theta amplitude increased and was comparable to shams during periods of comparable theta power. Conversely, pyramidal cells from both sham and injured animals were weakly entrained to theta overall. However, pyramidal cells from injured animals exhibited increased entrainment strength to theta with increasing theta amplitudes which resulted in significantly stronger entrainment at comparable theta power. During awake immobility, SWRs were reliably generated in both sham and injured animals, but ripple amplitudes were drastically reduced in injured animals. Together, these results demonstrate an overall desynchronization in the hippocampus following TBI, with reduced power/PAC and changes in neuronal entrainment occurring during the theta state while animals actively explored their environments, and reduced ripple amplitudes occurring in the non-theta state during quiet immobility. These results reveal specific TBI-associated pathophysiology across distinct network states which implicate multiple independent mechanisms that are thought to support learning and memory computations in the hippocampus. Specifically, deficits in the theta state may disrupt online encoding and retrieval of spatial information while deficits in the non-theta state may disrupt memory consolidation and planning during spatial navigation. These distinct mechanisms may differentially impact behavior, but likely all contribute to cognitive dysfunction following TBI.

### Layer Specific Changes

A loss of hippocampal oscillatory power in awake animals following TBI has previously been reported^40,42,45^, however, these studies used either monopolar electrodes or twisted wires which are difficult to localize to specific lamina. Localization of electrodes within lamina is important when examining oscillations, as power varies greatly across layers. The use of high-density laminar probes in this study allowed us to precisely localize electrodes to both *st. pyr* and *st. rad* in CA1 revealing a loss of oscillatory power in injured animals from both layers that was more pronounced in *st. pyr*, and a dramatic reduction in PAC in injured rats that was specific to *st. pyr*. We focus on theta and gamma because they are well characterized and have known roles in hippocampal processing, but it is important to note that injured animals showed reductions in power that span a broad range of frequencies as previously reported^45^. When this broadband shift was corrected for, theta power was still significantly lower in injured animals compared to shams, but the decrease in gamma power no longer reached statistical significance. Importantly, the broadband decrease in power was still limited to physiologically relevant frequencies (4-90 Hz) which may be due to a global reduction in or desynchronization of synaptic input to CA1, though the exact cause of this shift is unknown. Overall, injured animals exhibited a loss of broadband power, corrected theta power, and theta-gamma PAC that was more pronounced in *st. pyr* suggesting TBI-associated disruptions to cells and circuits specifically involving this layer.

One cell type implicated in these disruptions is parvalbumin positive (PV^+^) basket cells, which provide strong peri-somatic inhibition to CA1 pyramidal cells that can tightly control the timing of pyramidal cell firing through shunting inhibition^95,96^. PV^+^ interneurons as a class are entrained to theta and gamma oscillations and can entrain pyramidal cells to these oscillations^74–76^. Because PV^+^ basket cells target the soma of pyramidal cells and strongly contribute to generating theta and gamma, the reduced entrainment of interneurons in injured animals would predict the observed loss of theta and gamma power and PAC in *st. pyr*. Furthermore, the strong correlation between theta amplitude and pyramidal cell entrainment to theta could be a result of the strong temporal control that PV^+^ basket cells have over pyramidal cell firing. As theta amplitude increases, interneurons drastically increase their entrainment strength, which could more strongly entrain pyramidal cells to theta oscillations, resulting in the hyper-entrainment seen in pyramidal cells from injured animals at higher levels of theta power. In the non-theta state, PV^+^ basket cells are strongly recruited to fire during SWRs and support ripple generation^65,97–99^. Thus, the decrease in ripple power during SWR events may also suggest PV^+^ basket cell dysfunction. While hippocampal PV^+^ interneuron loss and dysfunction has previously been reported in a variety of injury models^41,100–102^, it is likely that TBI also disrupts other interneuron populations, which could contribute to the observed deficits especially the loss of oscillatory power in *st. rad*. Future studies recording from specifically identified interneuron classes would be needed to address whether TBI preferentially disrupts distinct interneuron subpopulations, and phase-specific stimulation could determine their contribution to disrupted theta and gamma oscillations in injured animals.

### Loss of Afferent Input

Disruptions to afferent input from CA3 and EC to CA1 can also contribute to electrophysiological deficits observed following TBI. CA3 pyramidal neurons are prone to excitotoxicity and are susceptible to cell loss in the _L_FPI model, and human cases of TBI^43,103–106^. Additionally, axonal injury has been demonstrated in pathways connecting CA3 and EC to CA1, which deafferents CA1, and may indicate neurodegeneration in these upstream regions^43,107^. Axonal injury is also known to cause action potential conduction delays and may therefore lead to desynchronization of CA1 afferent input^108,109^, which may be particularly important as rhythmic inputs from EC and CA3 are known to contribute to layer specific oscillations recorded in CA1. Concurrent recordings have not been performed between EC or CA3 and CA1 in awake, injured animals, so it is unknown whether rhythmic firing of cells in these regions is disrupted following TBI. Deafferentation of CA1 may also contribute to circuit hyperexcitability in TBI, which has been reported in both DG and CA1 in brain slices across different TBI models^25,31,33,34^. Compensatory mechanisms following TBI and/or deafferentation are known to alter the intrinsic properties of cells in CA1 and also disrupt long term potentiation^16,24,27–30,32,33^. Further, TBI has been shown to disrupt hyperpolarization-activated cation nonselective (HCN) channels responsible for *I*_h_ ^110–112^, which can affect cell excitability and could disrupt theta generation, particularly in the apical dendrites where these channels are expressed in large numbers. While homeostatic mechanisms may be responsible for the nonsignificant differences in firing rates between sham and injured animals, the increased recruitment of pyramidal cells across environments supports overall circuit hyperexcitability in CA1. The mechanism of this change is unknown but could be due to changes in inhibition, intrinsic excitability, or synaptic strength and efficacy following TBI. The interneuron specific changes we report along with the finding that pyramidal cell bursting was not changed following TBI support hyperexcitability due to altered inhibition, but we cannot rule out intrinsic or synaptic mechanisms.

TBI-associated changes in oscillatory power, PAC, and entrainment may also arise from desynchronization between the hippocampus and medial septum and diagonal band of Broca (MSDB). Reciprocal connections between MSDB and hippocampus lie within the fimbria and fornix where axonal pathology is reliably seen in the _L_FPI model^43,113^. Importantly, the extent of fimbria/fornix disruption has been correlated with poor learning and memory performance in human TBI^10–12^. In rodents, MSDB is especially important for theta generation and sets the pace of theta through theta frequency firing of MSDB interneurons^51,114–116^. Using the _L_FPI model, we previously demonstrated hippocampal-MSDB desynchronization in injured rats under isoflurane anesthesia^43^. Here, in awake rats, we show that CA1 interneurons are less entrained to theta in injured animals, which may be due to hippocampal-MSDB desynchronization as the entrainment of CA1 interneurons can be controlled by MSDB projecting interneurons^51,117^. Because CA1 interneurons are directly implicated in theta generation, the linear relationship between theta amplitude and entrainment strength suggests that the TBI-associated reduction in theta entrainment may be driving the overall loss of theta power observed in injured animals. Future studies performing simultaneous recordings from MSDB and hippocampus could help to dissociate whether these changes are driven by disrupted rhythmic inhibition from MSDB following TBI. MSDB glutamatergic and cholinergic inputs are also affected by fimbria/fornix pathology, which may further contribute to theta dysfunction in TBI. MSDB acetylcholine neurons have been demonstrated to be susceptible to injury resulting in cholinergic dysfunction^113,118,119^. This can disrupt type II theta, which occurs when animals are not actively moving and exploring the environment^51,120–122^. We found that single unit entrainment to theta was substantially decreased in injured rats when they were not moving, a phenomenon not seen in shams, which may suggest a disruption in type II theta. While these changes could be due to differences in attention and arousal or to the overall reduction in theta power at rest, our results suggest that cholinergic signaling may be dysfunctional following TBI.

### Implications of Temporal Coding Deficits

Oscillations provide the temporal framework to coordinate the firing of distinct cell ensembles within and across brain regions^47,123–125^. Theta oscillations arise from complex interactions between distributed circuits, which can temporally link distant brain regions in time, while gamma oscillations reflect processing important for selecting behaviorally relevant cell ensembles locally^50,53,54,123,125^. Thus theta-gamma PAC represents a mechanism whereby local processing in many distributed networks can be temporally coordinated on a theta timescale^47,59,126^. Importantly, LTP and LTD may preferentially occur at specific phases of the theta oscillation creating temporal windows for plasticity and potentially supporting spike-timing dependent plasticity across long distances^48^. In CA1, theta oscillations can provide temporal windows in which CA3 and EC inputs are integrated^47,127^. Additionally, coordinated CA3 and EC inputs recruit subsets of CA1 interneurons that drive layer-specific gamma oscillations^47,59,126^. CA3 driven low gamma (gamma_L_; 30-59 Hz) and EC driven mid gamma (gamma_M_; 60-80Hz) are respectively coupled to the descending phase (∼90°) and trough (∼180°) of theta measured in *st. pyr*^126^. This provides temporal separation of CA3 and EC inputs, which is hypothesized to prevent interference of previously stored associations in the CA3 recurrent network with the formation of new associations driven by EC^48^. Our findings of reduced theta-gamma PAC in TBI rats and a shift in the peak of gamma_L_ amplitude to a later phase of theta suggest that memory encoding and retrieval may not be sufficiently temporally separated in TBI rats. Further supporting this, we found that a larger percentage of pyramidal cells in injured animals fired across the two distinct environments, outlining a potential population coding deficit whereby a greater overlap in cell ensembles encoding these environments could result in less distinct neural representations. One proposed function of gamma is to select only relevant cell ensembles while inhibiting activity from other ensembles^123,125^. The loss of gamma power and theta-gamma PAC along with the over recruitment of pyramidal cells across environments suggests that this mechanism is dysfunctional in TBI, which could lead to interference between the encoding and retrieval of memories and disrupt memory updating.

Since information in the hippocampus is thought to be chunked on a theta timescale^38,78,79,81,126,127^, findings from this study would predict a disruption in theta phase precession and the formation of theta sequences, processes that require precise temporal integration of afferent inputs with local circuits^38,77–81^. This is further supported by our finding that pyramidal cells from both sham and injured animals fire flexibly across the theta cycle overall, but during periods of high theta power, pyramidal cells from injured animals significantly increase their entrainment strength and fire more rigidly relative to the theta phase than sham animals. This decreased flexibility during periods of high theta power may impair the formation of behaviorally relevant sequences and could contribute to cognitive deficits in TBI. We were unable to assess this because limited occupancy of the largely open environments prevented rigorous extraction and comparison of place cells and because recordings were not obtained while rats performed a measurable spatial memory task. However, previous studies have reported TBI-associated disruptions in place cell specificity and stability^5,41^. Thus, future studies tying dysfunctional place fields to deficits in phase precession and theta sequences during a spatial navigation task are needed to probe these mechanisms. In the non-theta state, SWRs are frequently observed during which cell ensembles are activated in the same or reverse temporal order they fired during behavior but on a compressed timescale^67,68^. This replay of activity is important for memory consolidation, recall, and updating processes as well as planning behaviors^65,66^. Here we showed that ripples had decreased amplitudes in TBI animals, which may impair these processes, but future studies are needed to address whether the replay of behaviorally relevant cells ensembles is affected by TBI.

The implications of these findings on learning and memory should be interpreted cautiously because recordings were not obtained while animals performed a spatial navigation task with a quantifiable outcome. However, we replicated a well-known spatial memory deficit in the MWM at the same injury level used for animals that underwent electrophysiological recordings and we found no differences in the swim speed or distance traveled in the MWM between sham and injured animals. Previous studies have also demonstrated that sham and injured rats perform comparably in a version of the MWM where the platform is visible or when a constant start location is used^90,128^. Together, this suggests that spatial coding deficits rather than potential sensory/motor deficits underlie impaired performance in the MWM following TBI implicating our electrophysiological findings in spatial learning and memory.

### Future Therapies

Neuromodulation therapies, including theta stimulation of the MSDB, fornix, or hippocampus, have been shown to improve learning and memory deficits in the _L_FPI model of TBI^129–132^. While the physiological mechanisms underlying behavioral improvements have not been investigated, results from this study provide outcome measures to target and examine with future therapies. For example, stimulation paradigms that successfully re-entrain interneurons to theta and gamma and boost oscillatory power and theta-gamma PAC while maintaining flexible firing of pyramidal cells may be more successful at restoring TBI-associated cognitive deficits. There is a large parameter space to explore when performing brain stimulation, so having electrophysiological targets may allow for faster optimization of stimulation parameters. Transplantation of stem cells or interneurons have also been shown to restore cognitive deficits in preclinical models of TBI^133,134^. Understanding how these transplanted cells functionally integrate into the network may help to unveil how pathophysiology correlates to behavior, especially if transplanted interneurons become entrained to local theta and gamma oscillations or if transplantation alters oscillatory power and coupling or pyramidal cell recruitment and spike timing.

Overall, TBI pathologies can vary across patients and have a wide range of effects on the brain. Fixing a single TBI-associated pathological change is unlikely to improve cognitive outcomes, thus it is important to study how coalescing TBI pathological changes disrupt neural circuit activity underlying cognition in order to develop better targets and treatments^70,135,136^. While some of the TBI pathophysiology we report here may be model and timepoint specific, similar electrophysiological deficits have been reported in Alzheimer’s disease, aging, epilepsy, and schizophrenia^137–146^, all of which can be associated with impaired cognition. Thus, temporal coding disruptions described in this study may be a good readout of hippocampal circuit function and could be predictive of cognitive dysfunction. As we learn more about how coding in the human hippocampus supports learning and memory, and how TBI disrupts these processes, we may be able to optimize deep brain stimulation treatments for TBI patients^147–150^ and potentially incorporate non-invasive tools such as transcranial magnetic stimulation^151–153^, transcranial alternating current stimulation^154,155^, and focused ultrasound^156,157^ to address cognitive impairments of TBI.

---

**Supplementary Figure 1:**
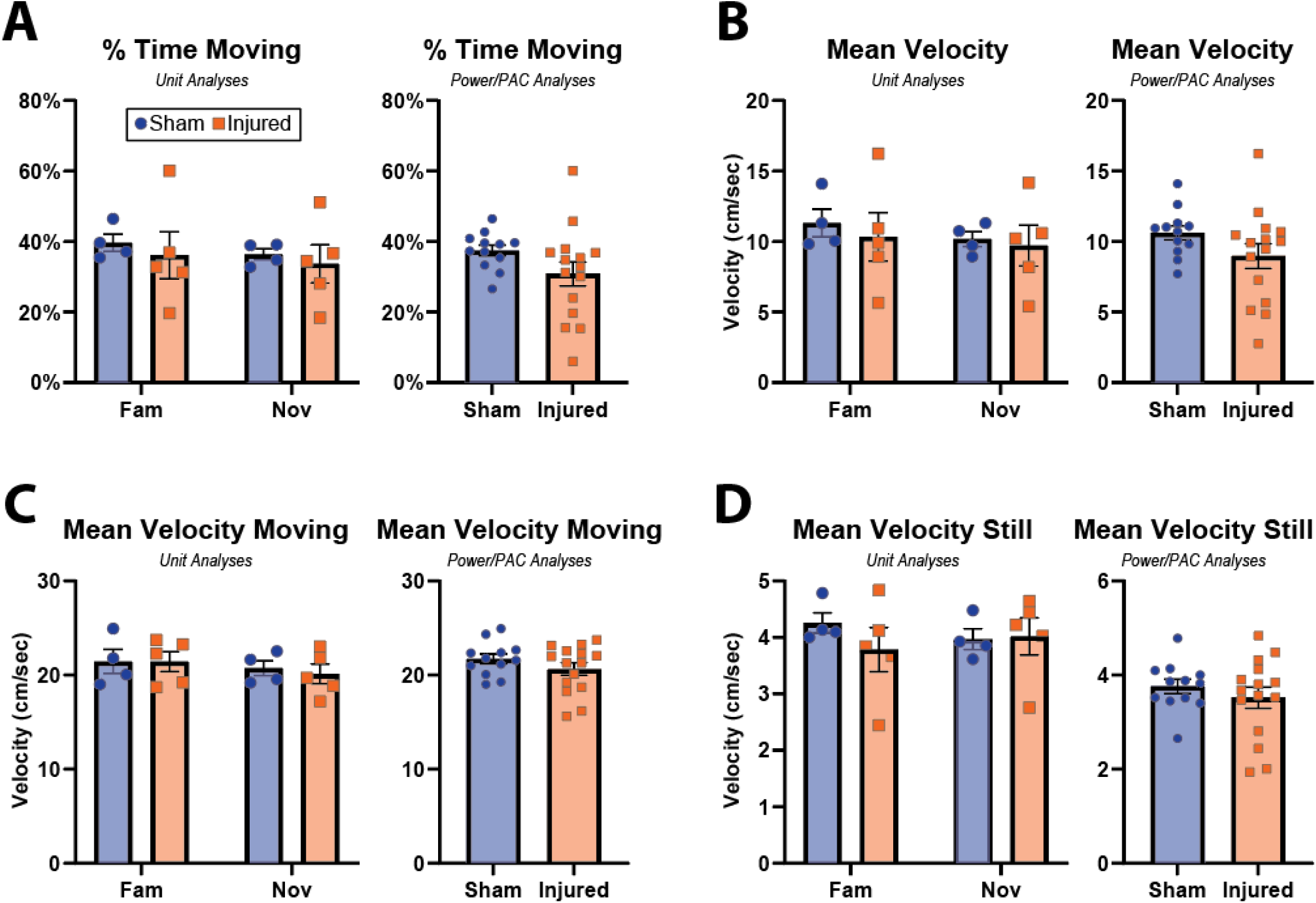
_L_FPI does not impact locomotion. **(A)** Left, percentage of time moving across groups and environments for recordings used for unit analyses (Sham Fam=39.66±2.42, Injured Fam=36.13±6.62, Sham Nov=36.39±1.57, Injured Nov=33.72±5.38). 2-way ANOVA reveals no significant effects of injury (p=0.667, f=0.202, df=1, 2.62% of total variation) or environment (p=0.088, f=3.766 df=1, 2.15% of total variation) on the percentage of time moving. Right, percentage of time moving in recordings used for power and PAC analyses (Sham=37.43±1.54, Injured=30.82±3.45; p=0.096, t-test with Welch’s correction). **(B)** Left, mean movement velocity (cm/sec) during recordings used for unit analyses (Sham Fam=11.32±0.98, Injured Fam=10.33±1.72, Sham Nov=10.18±0.53, Injured Nov=9.71±1.44). 2-way ANOVA reveals no significant effects of injury (p=0.709, f=0.1516, df=1, 1.948% of total variation, main effects model) or environment (p=0.084, f=3.903, df=1, 2.657% of total variation) on movement velocity. Right, mean movement velocity during recordings used for power and PAC analyses (Sham=10.61±0.49, Injured=8.97±0.87; p=0.114, t-test with Welch’s correction). **(C)** Left, mean velocity (cm/sec) during periods of movement in recordings used for unit analyses (Sham Fam=21.45±1.30, Injured Fam=21.44±1.04, Sham Nov=20.74±0.81, Injured Nov=20.12±1.04). 2-way ANOVA reveals no significant effects of injury (p=0.833, f=0.04786, df=1, 0.5603% of total variation) or environment (p=0.063, f=4.654, df=1, 6.43% of total variation) on movement velocity. Right, mean velocity during periods of movement in recordings used for unit analyses (Sham=21.71±0.52, Injured=20.65±0.66; p=0.234, t-test). **(D)** Left, mean velocity during periods when animals were not locomoting in recordings used for unit analyses (Sham Fam=4.26±0.18, Injured Fam=3.79±0.39, Sham Nov=3.97±0.18, Injured Nov=4.02±0.33). 2-way ANOVA reveals no significant effects of injury (p=0.641, f=0.2377, df=1, 3.036% of total variation) or environment (p=0.993, f=7.204e-5, df=1, 6.8e-5% of total variation) on movement velocity. Right, mean velocity during periods when animals were not locomoting in recordings used for power and PAC analyses (Sham=3.76±0.15, Injured=3.52±0.22; p=0.400, t-test). All comparisons utilized an ANOVA main effects model. For unit analyses, only day 1 of recording was used (n=4 and n=5 sessions for sham and injured respectively in each environment). For power and PAC analyses, all 3 days in the familiar environment were used (n=12 and n=15 sessions for sham and injured respectively in each environment).

**Supplementary Figure 2:**
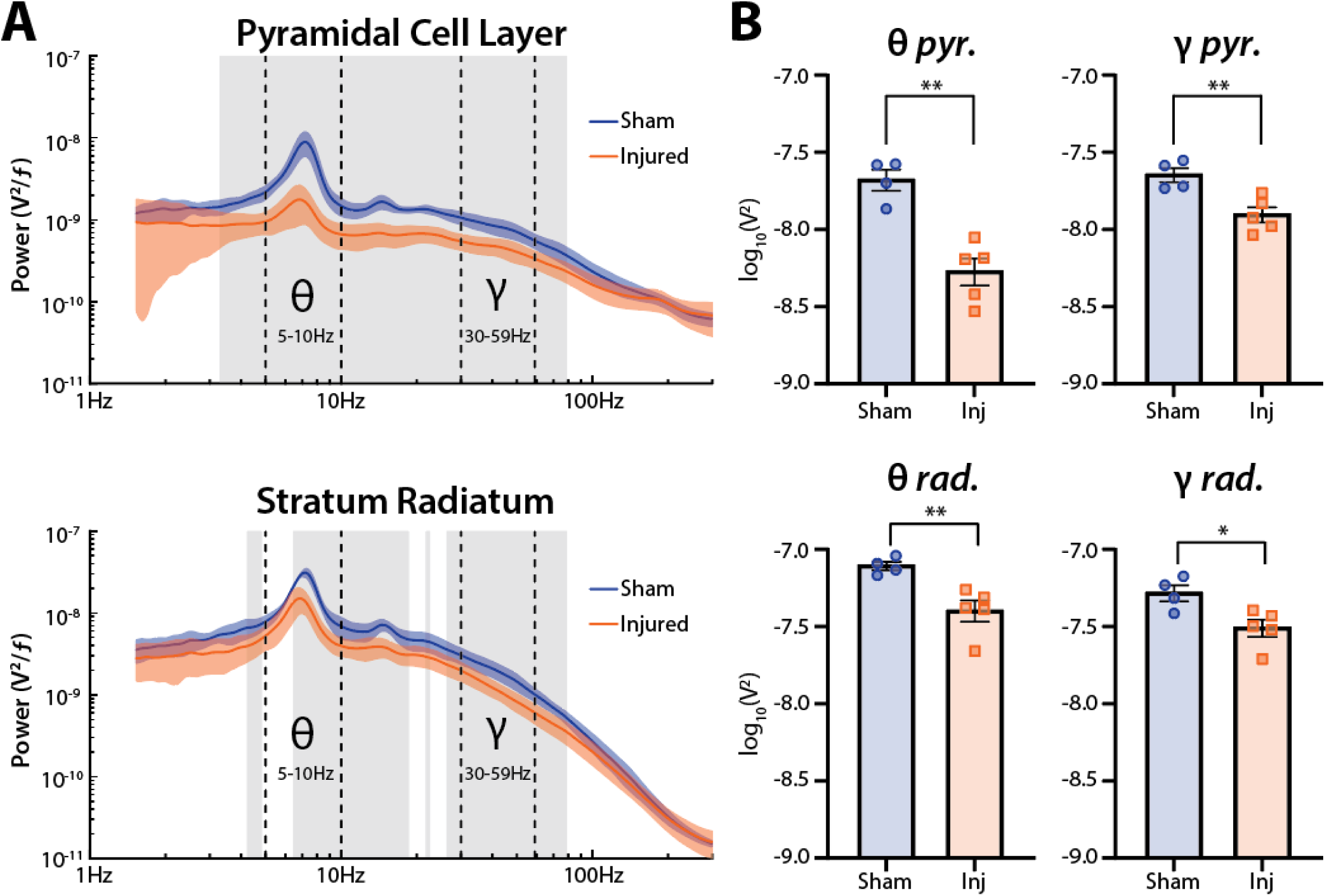
Power while still. **(A)** Power spectra (mean±SEM) from *st. pyr* (top) and *st. rad* (bottom) for sham and injured rats while not moving (<10 cm/sec) in the familiar environment. Theta (5-10 Hz) and gamma (30-59 Hz) bands denoted by dashed lines. Statistically significant differences between sham and injured are outlined in gray (α=0.01 for multiple comparisons; t-tests). **(B)** Integral of power in the theta and gamma frequency bands (mean±SEM; individual animals labeled by points) from sham and injured rats in *st. pyr* (top; theta: sham=-7.68±0.07 log_10_V^2^, injured=-8.28±0.09 log_10_V^2^, p=0.001; gamma: sham=-7.65±0.05 log_10_V^2^, injured=-7.91±0.05 log_10_V^2^, p=0.007; t-test) and *st. rad* (bottom; theta: sham=-7.11±0.03 log_10_V^2^, injured=-7.40±0.07 log_10_V^2^, p=0.009; gamma: sham=-7.28±0.05 log_10_V^2^, injured=-7.51±0.05 log_10_V^2^, p=0.022; t-test).

**Supplementary Figure 3:**
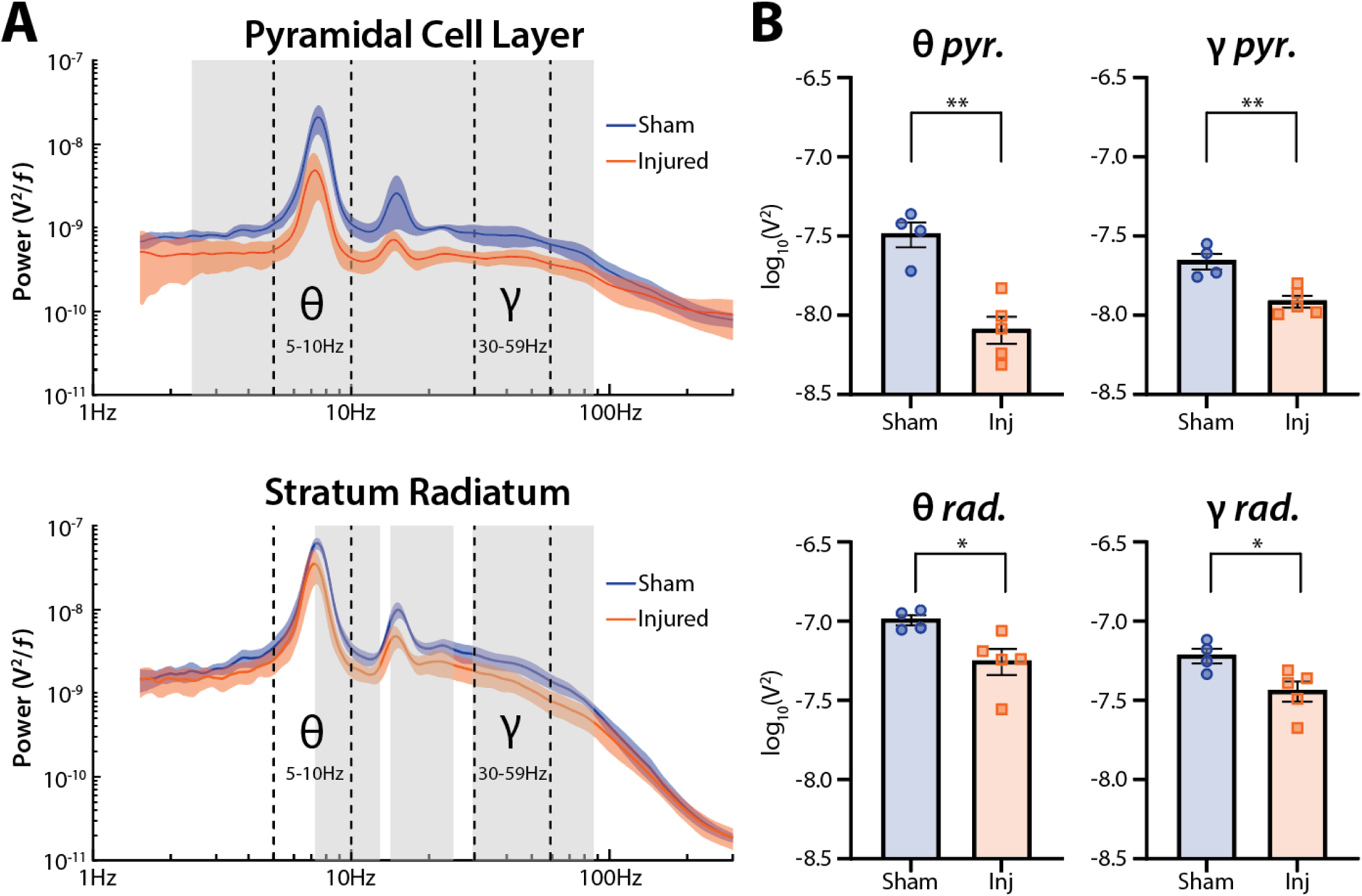
Power at higher movement velocities. **(A)** Power spectra (mean±SEM) from *st. pyr* (top) and *st. rad* (bottom) for sham and injured rats moving (>20 cm/sec) in the familiar environment. Theta (5-10 Hz) and gamma (30-59 Hz) bands denoted by dashed lines. Statistically significant differences between sham and injured are outlined in gray (α=0.01 for multiple comparisons; t-tests). **(B)** Integral of power in the theta and gamma frequency bands (mean±SEM; individual animals labeled by points) from sham and injured rats in *st. pyr* (top; theta: sham=-7.49±0.08 log_10_V^2^, injured=-8.10±0.09 log_10_V^2^, p=0.002; gamma: sham=-7.66±0.05 log_10_V^2^, injured=-7.92±0.04 log_10_V^2^, p=0.004; t-test) and *st. rad* (bottom; theta: sham=-6.99±0.03 log_10_V^2^, injured=-7.26±0.08 log_10_V^2^, p=0.030; gamma: sham=-7.22±0.05 log_10_V^2^, injured=-7.45±0.06 log_10_V^2^, p=0.032; t-test).

**Supplementary Figure 4:**
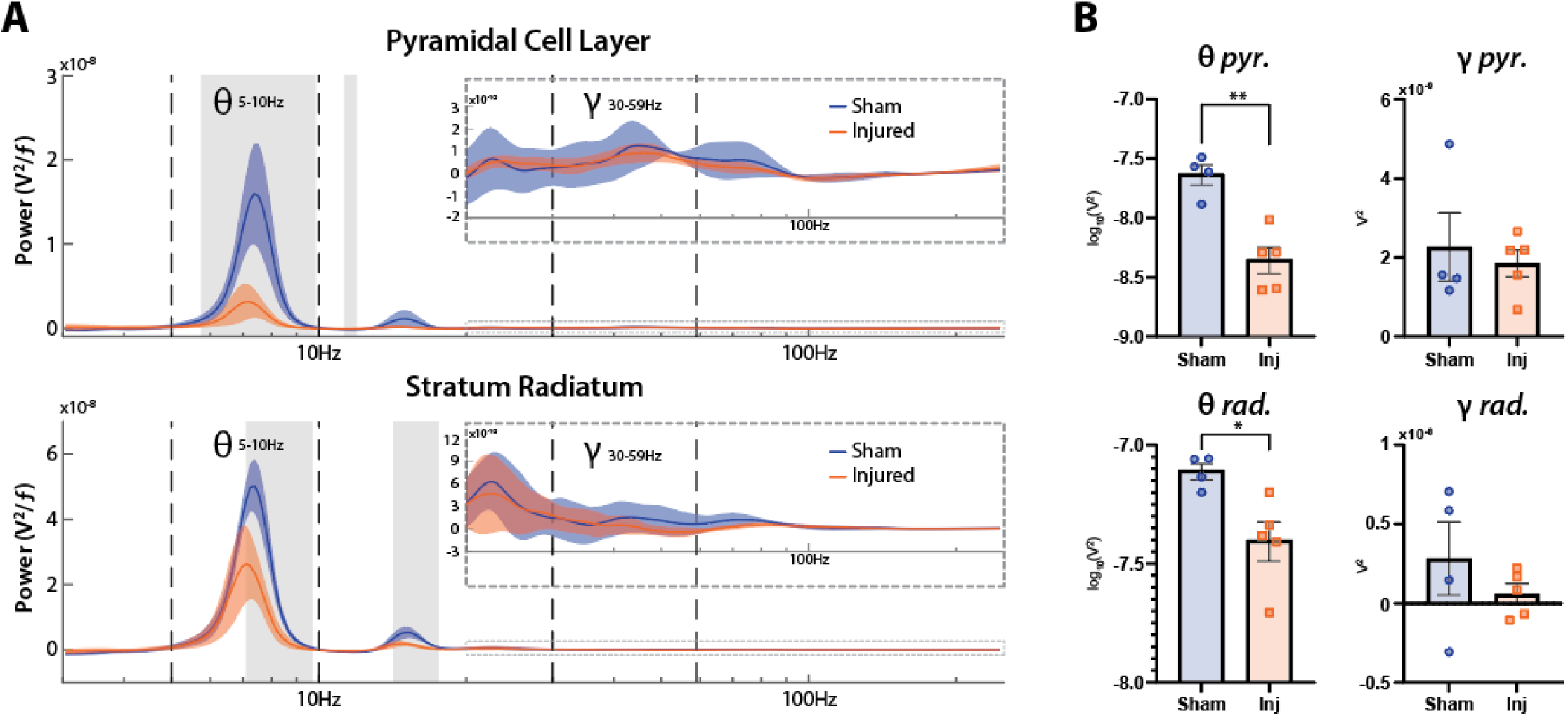
Broadband corrected power spectra. **(A)** Flattened power spectra (mean±SEM) from st. pyr (top) and st. rad (bottom) for sham and injured rats moving (>10 cm/sec) in the familiar environment. Theta (5-10 Hz) and gamma (30-59 Hz) bands denoted by dashed lines. Statistically significant differences between sham and injured are outlined in gray (α=0.01 for multiple comparisons; t-tests). Inset: zoomed in spectra in the higher frequency bands (>20Hz). **(B)** Integral of the corrected power spectra in the theta and gamma frequency bands (mean±SEM; individual animals labeled by points) from sham and injured rats in *st. pyr* (top; theta: sham=-7.64±0.09 log_10_V^2^, injured=-8.36±0.11 log_10_V^2^, p=0.002; gamma: sham=2.27x10^-9^±8.71x10^-10^ V^2^, injured=1.86x10^-9^±3.40x10^-10^ V^2^, p=0.645; t-test) and *st. rad* (bottom; theta: sham=-7.11±0.03 log_10_V^2^, injured=-7.41±0.08 log_10_V^2^, p=0.021; gamma: sham=2.84x10^-9^±2.31x10^-9^ V^2^, injured=6.05x10^-10^±6.44x10^-10^ V^2^, p=0.335; t-test).

**Supplementary Figure 5:**
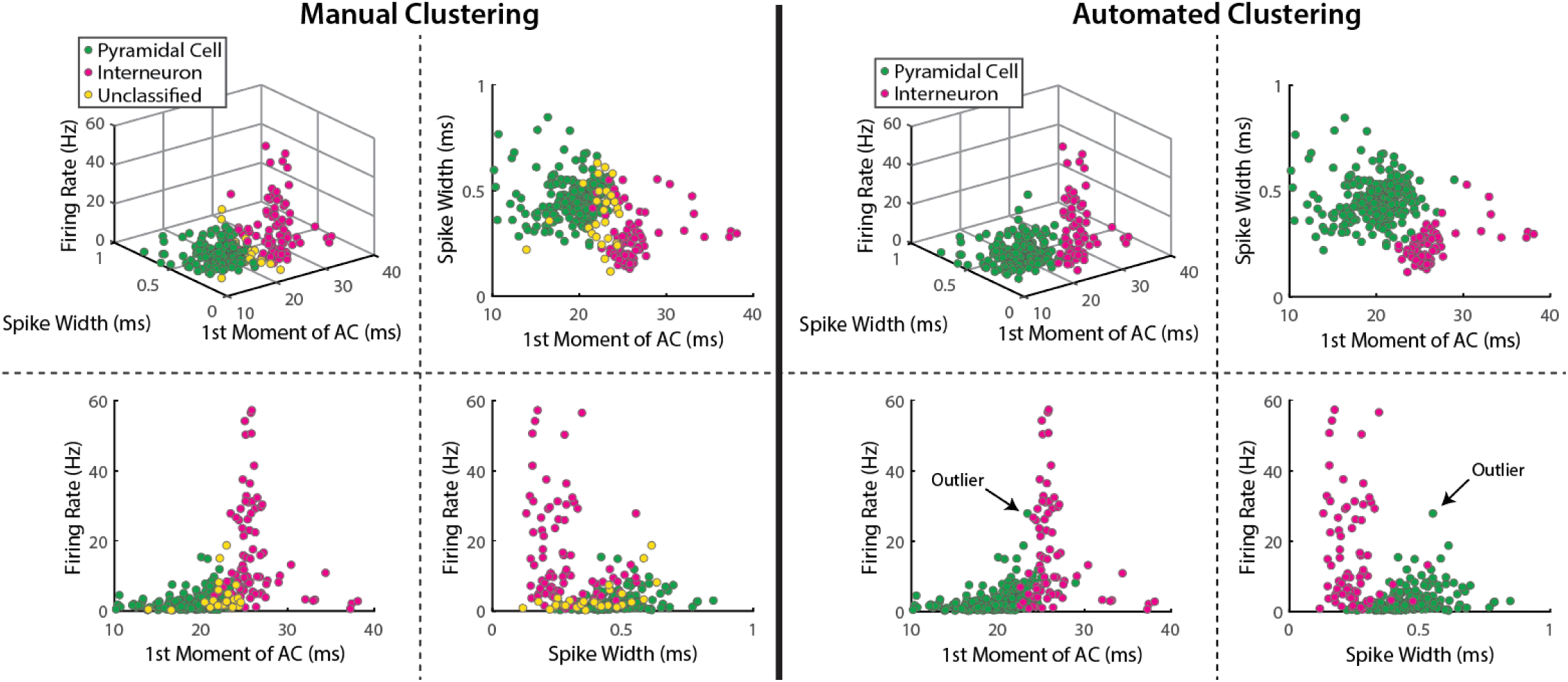
Comparison of manual and automated clustering of cell types. Scatter plots of spike width, first moment of the autocorrelogram, and firing rate for all single units located within the defined pyramidal cell layer. Left side shows manual clustering (matching **Fig 3B**) used for all analyses. Right side shows automated clustering (k-means consensus clustering with 2 groups). There was a 95.47% agreement between the two methods when unclassified cells were not included in the comparison. We chose to use manual clustering because inclusion of the unclassified group allowed us to be more conservative, and manual clustering is more robust to outliers in automated clustering such as the cell classified as a pyramidal cell with a firing rate of ∼30 Hz.

**Supplementary Figure 6:**
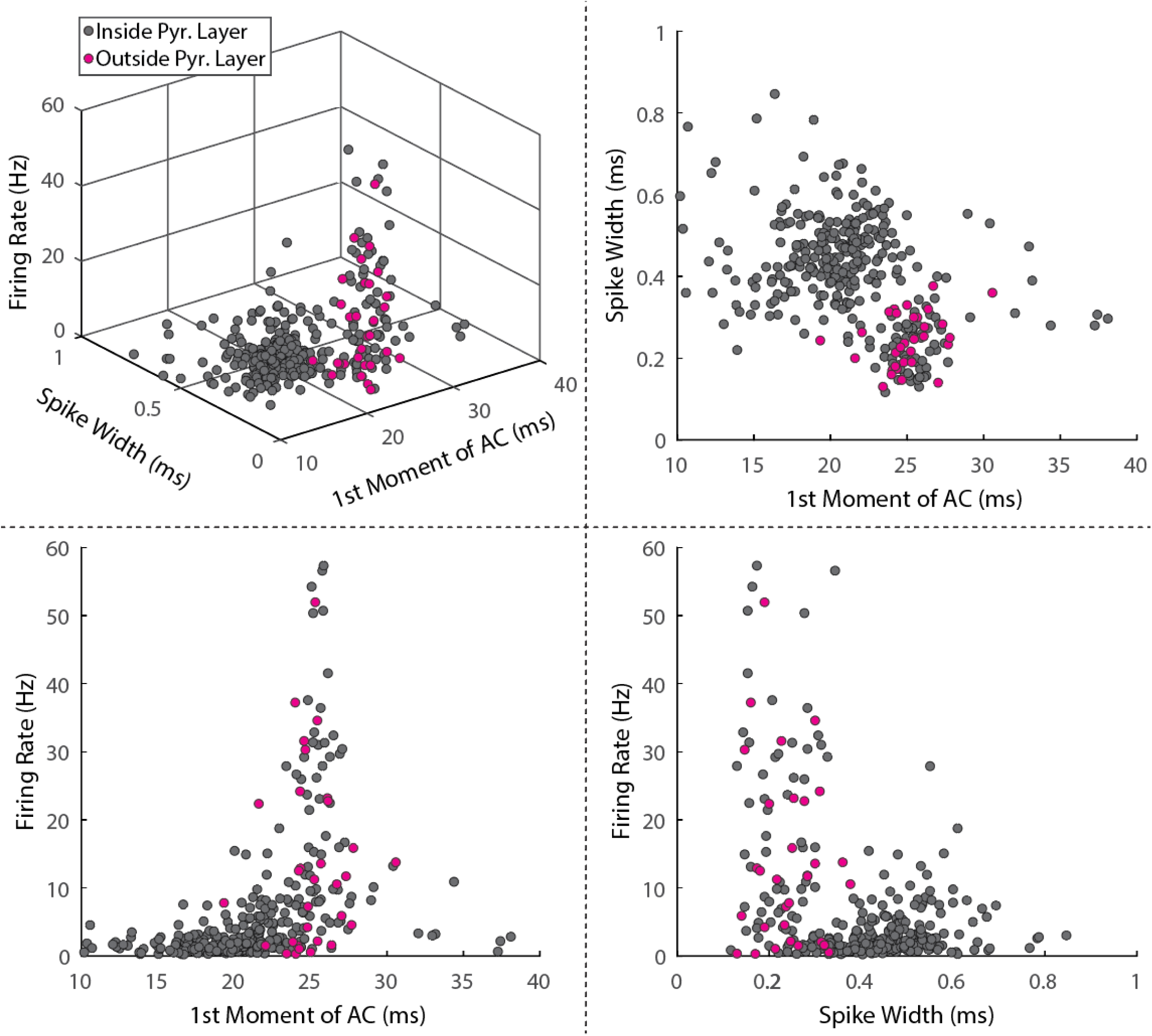
Cells above the defined pyramidal cell layer have features similar to interneurons. Scatter plots of spike width, first moment of the autocorrelogram, and firing rate for all single units. Cells in purple were >80 µm above the defined *st. pyr* channel and were automatically identified as interneurons. These cells have firing properties matching interneurons and cluster around the interneuron group (see Fig 3).

**Supplementary Figure 7:**
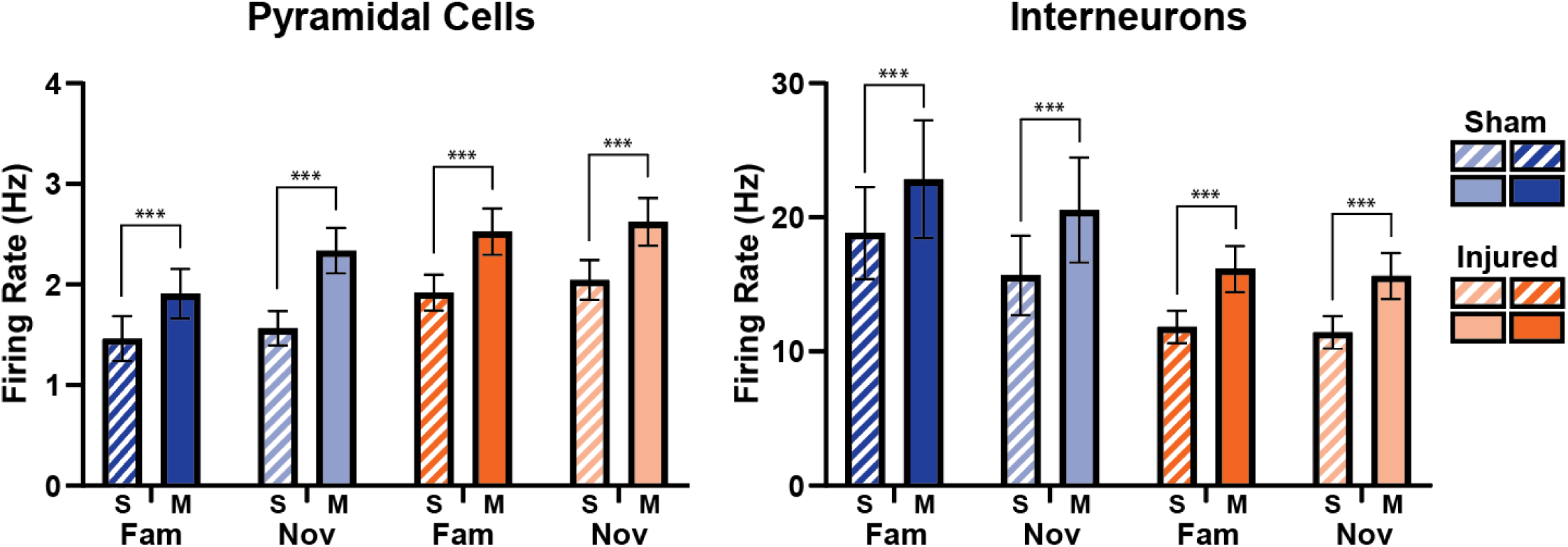
Firing rates while still or moving. Firing rates (Hz) across environments while animals were still (S) or moving (M; *Pyramidal cells:* Sham Fam. Still=1.46±0.22, Sham Fam. Moving=1.91±0.25, Sham Nov. Still=1.56±0.17, Sham Nov. Moving=2.34±0.22, Injured Fam. Still=1.92±0.18, Injured Fam. Moving=2.53±0.23, Injured Nov. Still=2.04±0.20, Injured Nov. Moving=2.62±0.24; *Interneurons:* Sham Fam. Still=18.83±3.42, Sham Fam. Moving=22.85±4.39, Sham Nov. Still=15.68±2.97, Sham Nov. Moving=20.54±3.91, Injured Fam. Still=11.84±1.21, Injured Fam. Moving=16.15±1.71, Injured Nov. Still=11.43±1.20, Injured Nov. Moving=15.63±1.71; mean±SEM). Firing rates were higher when animals were moving compared to when they were still across all conditions (Wilcoxon matched-pairs signed rank tests, p<0.001 for all conditions). There were no significant differences in firing rates between cells in sham and injured animals across any condition (*Pyramidal cells:* Fam. Still, p=0.212; Fam. Moving, p=0.579; Nov. Still, p=0.635; Nov. Moving, p=0.107; *Interneurons:* Fam. Still, p=0.093; Fam. Moving, p=0.216; Nov. Still, p=0.116; Nov. Moving, p=0.284; ks-tests).

**Supplementary Figure 8:**
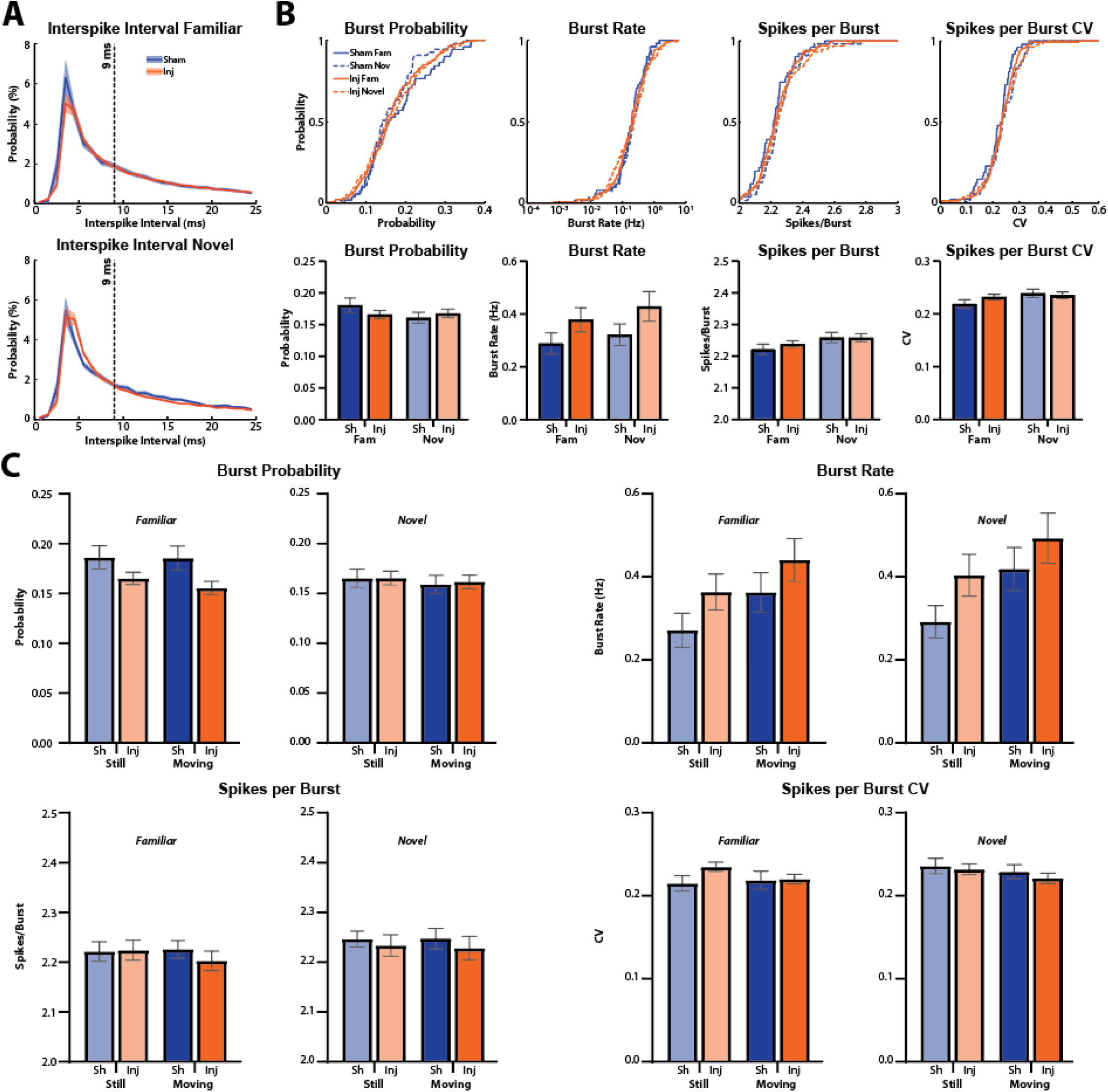
_L_FPI does not change pyramidal cell bursting. **(A)** Average interspike intervals across all pyramidal cells in the familiar and novel environment. No points along the curve were significantly different between sham and injured animals (ks-tests, α=0.01 for multiple comparisons). **(B)** Across all conditions, there were no significant differences in pyramidal cell burst probability (Sham Fam=0.18±0.01, Injured Fam=0.17±0.01, p=0.388; Sham Nov=0.16±0.01, Injured Nov=0.17±0.01, p=0.253), burst rate (Hz; Sham Fam=0.29±0.04, Injured Fam=0.38±0.05, p= 0.884; Sham Nov=0.32±0.04, Injured Nov=0.43±0.06, p= 0.561), spikes per burst (Sham Fam=2.22±0.02, Injured Fam=2.24±0.01, p=0.571; Sham Nov=2.26±0.02, Injured Nov=2.26±0.01, p=0.957), or spikes per burst coefficient of variation (Sham Fam=0.22±0.01, Injured Fam=0.23±0.01, p=0.467; Sham Nov=0.24±0.01, Injured Nov=0.24±0.01, p=0.998) between sham and injured animals (mean±SEM, ks-test used for all comparisons). **(C)** When further split into periods when animals were still or moving, there were no differences in burst probability (Familiar: Sham Still=0.19±0.01, Injured Still=0.17±0.01, p=0.401; Sham Moving=0.19±0.01, Injured Moving=0.16±0.01, p=0.145; Novel: Sham Still=0.17±0.01, Injured Still=0.17±0.01, p=0.667; Sham Moving=0.16±0.01, Injured Moving=0.16±0.01, p=0.287), burst rate (Familiar: Sham Still=0.27±0.04, Injured Still=0.36±0.04, p=0.610; Sham Moving=0.36±0.05, Injured Moving=0.44±0.05, p=0.674; Novel: Sham Still=0.29±0.04, Injured Still=0.40±0.05, p=0.406; Sham Moving=0.42±0.05, Injured Moving=0.49±0.06, p=0.238), spikes per burst (Familiar: Sham Still=2.22±0.02, Injured Still=2.22±0.02, p=0.357; Sham Moving=2.23±0.02, Injured Moving=2.20±0.02, p=0.330; Novel: Sham Still=2.25±0.02, Injured Still=2.23±0.02, p=0.953; Sham Moving=2.25±0.02, Injured Moving=2.23±0.02, p=0.406), or spikes per burst coefficient of variation (Familiar: Sham Still=0.21±0.01, Injured Still=0.24±0.01, p=0.100; Sham Moving=0.22±0.01, Injured Moving=0.22±0.01, p=0.749; Novel: Sham Still=0.24±0.01, Injured Still=0.23±0.01, p>0.999; Sham Moving=0.23±0.01, Injured Moving=0.22±0.01, p=0.874) across all conditions between sham and injured animals (mean±SEM, ks-test used for all comparisons).

**Supplementary Figure 9:**
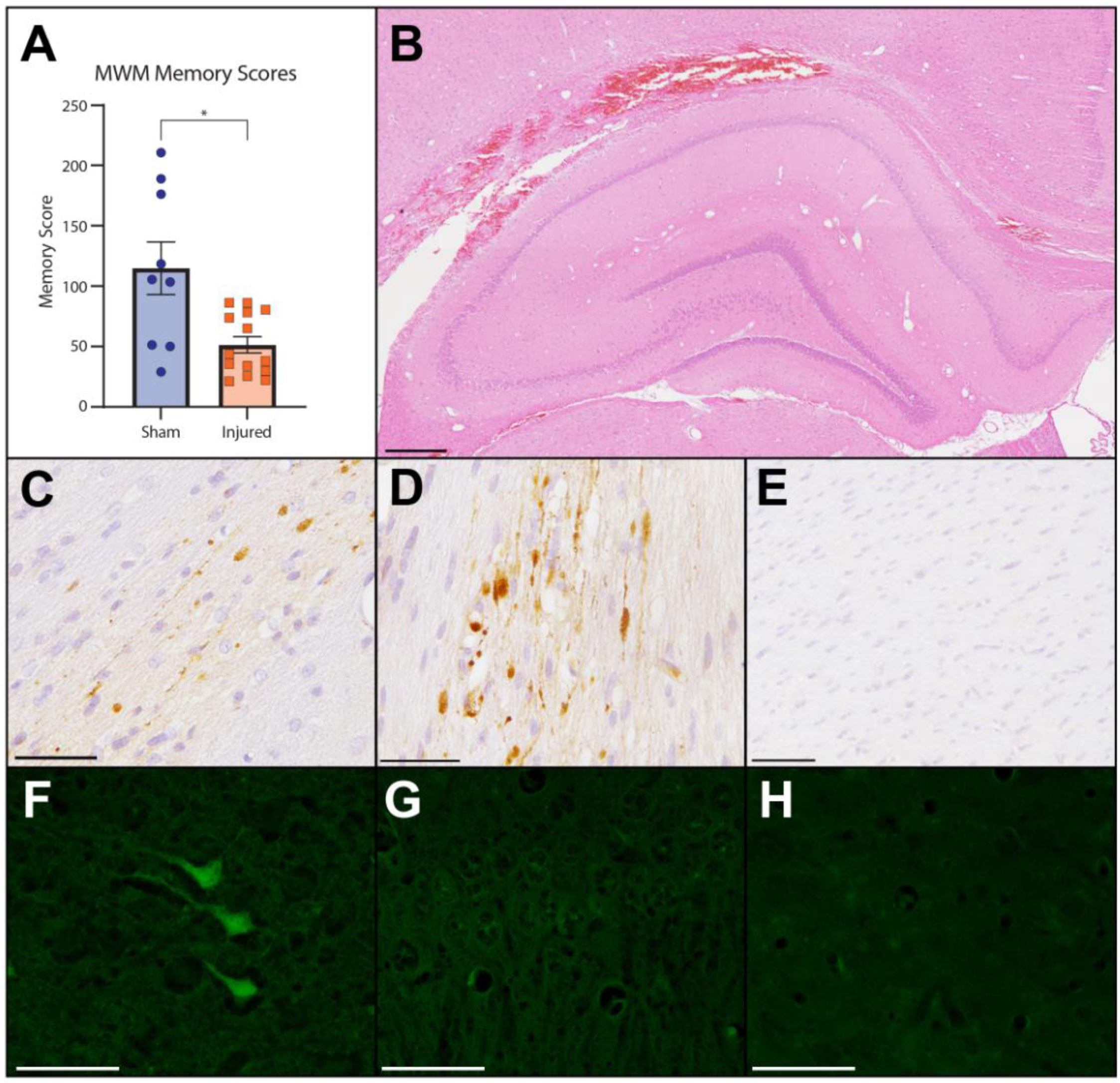
_L_FPI disrupts spatial memory and induces stereotypic pathologies. **(A)** Morris water maze memory scores (mean±SEM; individual animals labeled by points) from sham and injured rats tested at 48hr post-injury (sham=114.8±21.8, n=9, injured=51.5±6.8, n=14, p=0.020; Welch’s t-test). **(B)** H&E-stained sections showing hemorrhagic contusion in the ipsilateral white matter, including the corpus callosum at 48 hr post-_L_FPI. Note, the underlying hippocampus appears grossly intact (scale bar: 500 µm). **(C-D)** APP immunoreactive axonal pathology in the angular bundle (C) and fimbria-fornix (D) at 48 hrs post-_L_FPI (scale bars: 50 µm). **(E)** An absence of axonal pathology in the fimbria-fornix 48 hrs following sham procedures (scale bar: 100 µm). **(F)** Fluoro-Jade C positive neurons in the peri-lesional cortex at 48 hrs post-_L_FPI (scale bar: 50 µm). **(G)** An absence of Fluoro-Jade C positive cells in the CA1 region of hippocampus at 48 hrs post-_L_FPI (scale bar: 50 µm). **(H)** Ipsilateral cortex displaying an absence of Fluoro-Jade C positive cells following sham procedures (scale bar: 50 µm).

**Supplementary Figure 10:**
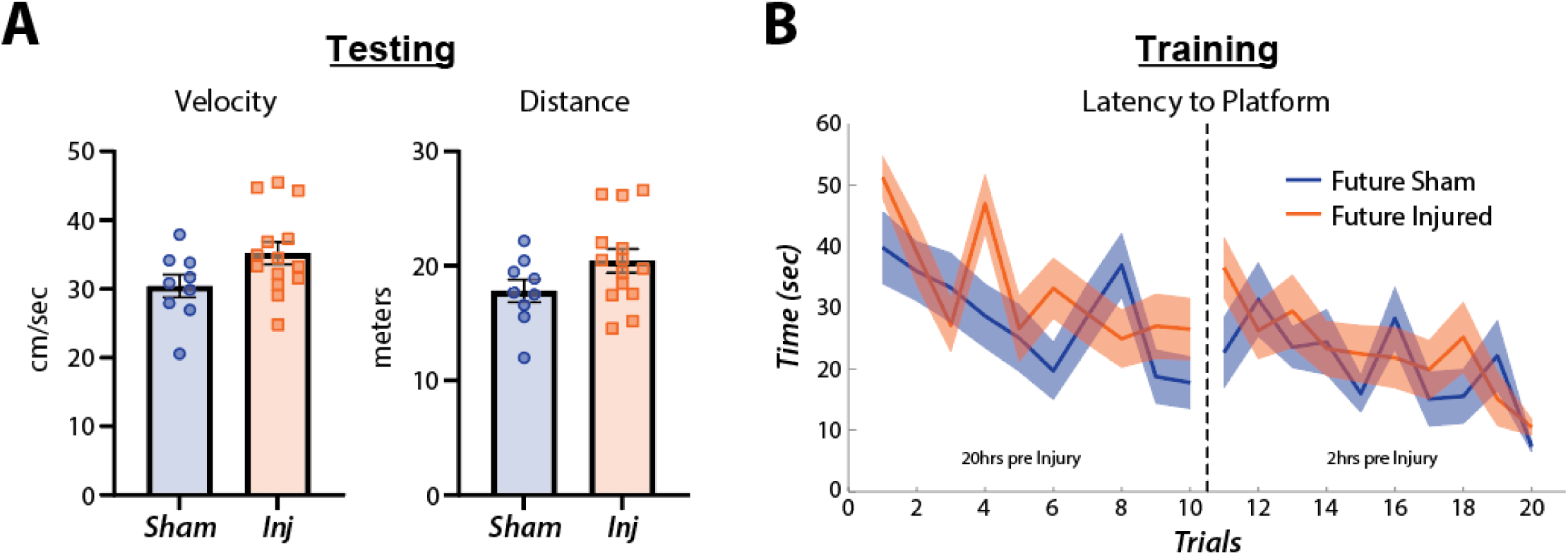
Additional MWM data. **(A)** There was no significant difference in swim velocity (mean±SEM; Sham=30.41±1.66, n=9; Injured=35.20±1.63, n=14; p=0.063, t-test) or distance traveled (Sham=17.82±0.99, n=9; Injured=20.44±1.03, n=14; p=0.098, t-test) during testing between sham and injured animals. **(B)** Learning curves indicate no differences in learning between the two groups prior to sham/_L_FPI surgery (2-way ANOVA showed a significant effect of learning across days p<0.001, f=2.929, df=19, 13.03% of total variation, but no effect across groups p=0.448, f=0.6079, df=1, 0.62% of total variation). Data is from a subset of n=7 sham and n=10 injured animals in which training sessions were recorded (they were not recorded in the other n=2 sham and n=4 injured).

**Supplementary Table 1:** Total number of isolated single units across rats.

| NUMBER OF CELLS ACROSS ANIMALS |  |  |  |
| --- | --- | --- | --- |
| Animal ID | Sham/Injured | Probe | Total Cells |
| Animal 1 | Sham | E1 (4 Shank) | 0* |
| Animal 2 | Sham | H2 (2 Shank) | 16 |
| Animal 3 | Injured | E1 (4 Shank) | 114 |
| Animal 4 | Injured | H2 (2 Shank) | 20 |
| Animal 5 | Injured | E1 (4 Shank) | 76 |
| Animal 6 | Injured | H2 (2 Shank) | 18 |
| Animal 7 | Sham | E1 (4 Shank) | 60 |
| Animal 8 | Sham | H2 (2 Shank) | 9 |
| Animal 9 | Injured | E1 (4 Shank) | 36 |
| *Electrode drift across sessions made rigorous comparisons not possible |  |  |  |

**Supplementary Table 2:** Additional statistics from t-tests.

| **t-test Comparisons** | **Location** | **p** | **t** | **df** | **tail** | **Note** |
| --- | --- | --- | --- | --- | --- | --- |
| Theta Power *pyr.* | Fig 1 G | 0.002 | 5.005 | 7 | 2 |  |
| Gamma Power *pyr.* | Fig 1 G | 0.007 | 3.803 | 7 | 2 |  |
| Theta Power *rad.* | Fig 1 G | 0.022 | 2.934 | 7 | 2 |  |
| Gamma Power *rad.* | Fig 1 G | 0.028 | 2.766 | 7 | 2 |  |
| PAC *pyr.* | Fig 2D | 0.006 | 3.945 | 7 | 2 |  |
| PAC *rad.* | Fig 2D | 0.623 | 0.514 | 7 | 2 |  |
| SWR Rate - Sham | Fig 6C | 0.011 | 5.658 | 3 | 2 |  |
| SWR Rate - Injured | Fig 6C | 0.052 | 2.735 | 4 | 2 |  |
| % Time Moving | Sup Fig 1A | 0.096 | 1.751 | 19.17 | 2 | Welch's correction |
| Mean Velocity | Sup Fig 1B | 0.114 | 1.647 | 21.55 | 2 | Welch's correction |
| Mean Velocity Still | Sup Fig 1C | 0.234 | 1.220 | 25 | 2 |  |
| Mean Velocity Moving | Sup Fig 1D | 0.400 | 0.856 | 25 | 2 |  |
| Theta Power *pyr.* | Sup Fig 2B | 0.001 | 5.163 | 7 | 2 |  |
| Gamma Power *pyr.* | Sup Fig 2B | 0.007 | 3.745 | 7 | 2 |  |
| Theta Power *rad.* | Sup Fig 2B | 0.009 | 3.574 | 7 | 2 |  |
| Gamma Power *rad.* | Sup Fig 2B | 0.022 | 2.935 | 7 | 2 |  |
| Theta Power *pyr.* | Sup Fig 3B | 0.002 | 5.037 | 7 | 2 |  |
| Gamma Power *pyr.* | Sup Fig 3B | 0.004 | 4.203 | 7 | 2 |  |
| Theta Power *rad.* | Sup Fig 3B | 0.030 | 2.725 | 7 | 2 |  |
| Gamma Power *rad.* | Sup Fig 3B | 0.032 | 2.682 | 7 | 2 |  |
| Theta Power *pyr.* | Sup Fig 4B | 0.002 | 4.935 | 7 | 2 |  |
| Gamma Power *pyr.* | Sup Fig 4B | 0.645 | 0.482 | 7 | 2 |  |
| Theta Power *rad.* | Sup Fig 4B | 0.021 | 2.976 | 7 | 2 |  |
| Gamma Power *rad.* | Sup Fig 4B | 0.335 | 0.104 | 7 | 2 |  |
| MWM Memory Score | Sup Fig 9A | 0.020 | 3.295 | 21 | 2 | Welch's correction |
| MWM Velocity | Sup Fig 10B | 0.063 | 1.965 | 21 | 2 |  |
| MWM Distance | Sup Fig 10B | 0.098 | 1.733 | 21 | 2 |  |

## METHODS

### Animals

All procedures and animal care related to this study were approved by the University of Pennsylvania Institutional Care and Use Committee at an Association for Assessment and Accreditation of Laboratory Animal Care (AAALAC) accredited site. Young healthy adult male Long Evans rats (RGD catalog: 2308852, Charles River Laboratories; RRID: RGD_2308852) naïve to any previous procedures or investigations were used in this study. Rats were kept on a strict 12 hr light/dark cycle and had access to standard chow and water *ad libitum* except when food restricted for electrophysiological experiments (see below). Animals were examined daily for signs of pain, or distress after all surgical procedures.

### _L_FPI Surgery

Lateral fluid percussion injury (_L_FPI) surgeries were performed as described previously^43^. Briefly, rats were anesthetized with isoflurane and affixed to a stereotaxic holder. A midline incision was made on the head to visualize the skull, then a 5mm craniectomy was made over the left parietal cortex centered between lambda and bregma in the anterior/posterior direction and between midline and the lateral edge of the skull in the medial/lateral direction. A needle hub (cut from the needle) was affixed to the craniectomy site and secured with glue and dental cement. Rats were then removed from anesthesia, monitored for increased respiration and a response to toe pinch, then immediately attached to the FPI device. To induce injury, a pendulum was dropped to strike a tube filled with sterile saline delivering a fluid pulse to the intact dura within the craniectomy. A transducer was used to measure the force of the pressure wave. The needle hub was then removed and Kwik-Sil was place over the site of the craniectomy. The incision was then sutured up and rats were placed under a warming lamp and administered buprenorphine (0.05 mg/kc s.c.) after demonstrating a righting reflex. Sham rats underwent the same surgical procedure as injured rats, including the craniectomy, but no fluid pulse was delivered. Animals were excluded from further study if a dural breach was present or if the craniectomy was not properly centered between lambda/bregma and midline/lateral edge. Four rats (3 injured and 1 sham) were excluded from the MWM based on these criteria.

### Electrode Implantation Surgery

Rats underwent electrode implantation surgery 3-4 days after _L_FPI/sham surgery. During this procedure, rats were anesthetized with isoflurane and affixed to a stereotaxic holder. Sutures from the FPI procedure were removed, the midline incision was re-opened to expose the skull surface, and the Kwik-Sil cover over the _L_FPI craniectomy was removed. A stainless-steel screw was placed on the surface of the dura over the cerebellum and was used as ground. A small tungsten wire (∼10 kohms) was placed in the lateral ventricle (0.7 mm posterior to bregma, 2.8 mm lateral (assuming angle), 3.8 mm DV, at a 10° angle lateral to medial) and was used as a reference. A durotomy was then performed over the _L_FPI craniectomy and electrodes mounted to drives (nano-Drive; Cambridge NeuroTech; Cambridge, United Kingdom) were slowly advanced into cortex (AP: -4.2 mm, ML: 2.8 mm, DV: ∼1.3 mm). Dura-Gel (Cambridge NeuroTech) was then applied to the brain surface and Vaseline was melted onto the electrode and cable using a low-temp cauterizer to allow them to move when advancing the electrode post-implant. Anchor screws were inserted bilaterally into the lateral ridge of the skull, and dental cement was applied to secure the implant. Rats were either implanted with a Cambridge NeuroTech E1 (4 shank) electrode (n=2 sham, n=2 injured) or a Cambridge NeuroTech H2 electrode (n=2 sham, n=3 injured).

### Behavior During Electrophysiological Recordings

Animals used for electrophysiological recordings were handled for 5-7 days then were food restricted (up to 85% of body weight) and introduced to a 1 m^2^ open field environment containing small pieces of Fruit Loops to encourage exploration. Rats were exposed to the environment for 15 mins/day for a minimum of 3 days before injury to become familiarized to it. Rats were then returned to normal *ad libitum* food access for at least 24 hrs then subjected to _L_FPI/sham surgery. Rats were chronically implanted with recording electrodes 3-4 days after sham/injury surgery. 24 hrs after implantation, rats were again food restricted (up to 85% of body weight) and habituated to the wireless recording transmitter (FreeLynx; Neuralynx; Bozeman, MT) while exploring the same open field environment for food rewards. The following days, electrodes were advanced deeper into the brain, and the rat was again placed in the open field environment to find food rewards. This process was repeated until electrodes reached *st. pyr* in CA1 (6-9 days post-injury) which was determined based on real-time observation of multi-unit activity and the presence of sharp-wave ripples. Once in *st. pyr*, rats were placed in their home cage for at least 20 mins to allow electrodes to settle then recordings were obtained in the familiar open field environment. Rats were then placed back in their home cage for 5 mins while the battery for the wireless transmitter was replaced, then were placed into a novel environment for an additional recording. The novel environment was an 8-arm radial arm maze with Fruit Loops scattered throughout to encourage exploration (no behavioral task was performed in the radial arm maze). On 2 subsequent days, electrodes were advanced further into the CA1 lamina, allowed to settle, then rats were recorded in the familiar open field environment.

### Data Acquisition

Electrical signals were sampled at 30 kHz, amplified, digitized, then wirelessly transmitted using the FreeLynx (Neuralynx). Real-time signals were visualized in the Cheetah recording software (Neuralynx) as local field potentials (LFPs) and filtered signals (600-6000 Hz) were organized into tetrodes and thresholded for real-time spike detection.

### Selection of St. Pyramidale and St. Radiatum Channels

On each shank, only a single *st. pyr* and *st. rad* channel was chosen. The channel containing the local maximum of power in the ripple frequency (100-250 Hz) range was chosen as the *st. pyr* channel. As electrodes were advanced on later recording days, *st. pyr* was not always present on every electrode shank thus only shanks with a local maximum of ripple frequency power were included. The *st. rad* channel was chosen as the local maximum of the CSD sink for the mean sharp-wave ripple (SWR) waveform. A local maximum of the SWR CSD sink was not observed on all shanks especially on the first recording day when most electrodes were localized to *st. pyr*, thus, shanks without a local maximum in the SWR CSD sink were not included.

### Velocity Calculations

The position of the animal was obtained by tracking a light on the wireless transmitter affixed to the rat’s head. Tracking was curated by manually removing artifacts (sudden large displacements), interpolating between removed positions, and smoothing pixel locations with a gaussian kernel. Pixels were then converted to real-world distances (cm) using the size of the environments for scaling. Instantaneous velocity (Δ position / Δ time) was computed between consecutive frames then smoothed with a gaussian kernel. For velocity comparisons in **Sup Fig 1**, movement velocity and the percentage of time rats spent locomoting (>10cm/sec) was evaluated to match other physiological comparisons. Unit analyses (firing rate and entrainment) were constrained to recording day 1 in each environment. Thus, movement was quantified in these epochs for each animal (left panel in **Sup Fig 1A-D**). Power and PAC analyses were constrained to familiar environment but included all 3 recording days. Thus, movement was quantified for each recording session (3/rat; right panel in **Sup Fig 1A-D**). Mean velocity was compared between sham and injured animals for the entire recording as well as epochs when animals were still (<10cm/sec) or moving (>10cm/sec).

### Power Analyses

Raw signals were down sampled to 3 kHz using an anti-aliasing filter (resample function in MATLAB; MathWorks, MA). Power was then computed using a continuous wavelet transform using the cwt function in MATLAB. First a filter bank of complex bump wavelets was created from 1-1500 Hz using 36 wavelets per octave. Bump wavelets were selected because they have narrow variance in frequency. Wavelets were then convolved with the down sampled signal and power was computed as the magnitude of the analytic signal squared for each convolution. For each recording in the familiar environment, the mean power for each frequency was computed specifically for times when the rat was moving (>10 cm/sec). These values were then averaged across shanks and days to get an average per animal for the defined *st. pyr* and *st. rad* channels respectively, and sham and injured animals were averaged with each other to create the power spectra shown in **Fig 1F**. Comparisons between power spectra of sham and injured animals were done using t-tests at each frequency in the plot. Significant differences outlined in grey were determined using an alpha of 0.01. Theta and gamma power was extracted from each animal’s mean power spectrum by computing the area under the curve in the 5-10 and 30-59 Hz bands respectively. These individual values are shown in **Fig 1G**. For broadband power corrections, data was first broken into 2 second epochs^45^, and a power spectrum was computed for each epoch. Only epochs in which the animal’s movement velocity was above the threshold of 10cm/sec for at least 50% of the time were included. We then modeled the aperiodic component of the power spectrum in each epoch (which can represent a broadband offset or change in exponent of the power spectra) using a Lorentzian function^69^, subtracted this component in each epoch, then averaged across epochs. We then averaged corrected spectra across animals, compared the spectra point-by-point between sham and injured animals as described above, and extracted theta and gamma bands as described for further comparison.

### Current Source Density

The current source density (CSD) was calculated across electrode channels on a shank using the spline inverse CSD method from^158^ in the freely available MATLAB CSDplotter (v0.1.1) package.

### PAC Analyses

The strength of theta-gamma PAC was assessed independently in *st. pyr* and *st. rad* by calculating the modulation index (MI)^71^ while rats were moving (>10 cm/sec) in the familiar environment. Data was down sampled to 600Hz, separately filtered into theta (5-10 Hz) and gamma (30-59Hz) frequency bands using a 2^nd^ order Butterworth filter, and the instantaneous theta phase and gamma amplitude was extracted for each sample using the Hilbert transform. The mean gamma amplitude (𝑎̅) was calculated across 18 theta phase bins (𝑘; 20° width) for epochs when the rat was moving then converted to a probability distribution (equation 1).

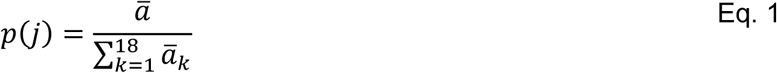

Shannon Entropy (𝐻) was calculated (equation 2) for the distribution, then the modulation index (𝑀𝐼) was computed (equation 3) and averaged across shanks and recording days to get a mean MI per rat (individual points in **Fig 2D**).

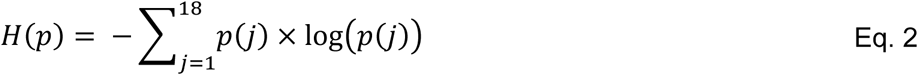

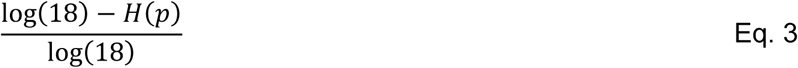

**Figure 2E** depicts the mean gamma amplitude across theta phase bins averaged across shanks and days to get a mean distribution for each rat, then averaged across animals in the sham and injured conditions. This distribution was then up sampled by a factor of 100 and the theta phase angle corresponding to the peak in gamma amplitude was extracted to compare between sham and injured animals (theta peak=0°). PAC heatmaps in **Fig 2B** were constructed by calculating the MI across a range of low-frequency bins for phase (40 logarithmically spaced bins spanning 1-15 Hz each with a width of 1 Hz) and high-frequency bins for amplitude (20 logarithmically spaced bins spanning 20-180 Hz each with a width of 20 Hz). Heatmaps were averaged across shanks and recording days to get an average for each rat, then averaged across sham and injured animals. For the theta amplitude vs PAC plot (**Fig 5E**), theta amplitude was split into 5 bins (see interneuron entrainment vs theta amplitude methods below for bin definitions) and PAC was calculated independently for each bin. A line was then fit to the data using linear regression.

### Single Unit Isolation and Clustering

Automated spike sorting was done using Klusta (RRID: SCR_014480) and manually curated in Phy (https://github.com/kwikteam/phy) by a reviewer blind to sham/injured condition. Each shank was assessed independently as the inter-shank spacing of 250 µm ensured that units were not visible on multiple shanks. Familiar and novel environment recordings on day 1 were spliced together, and spike sorting was performed on this joined data to retain unit identity across recordings. After single units were extracted, they were clustered into 3 different groups: putative pyramidal cells, interneurons, or unclassified. All units >80 µm above the defined *st. pyr* channel were automatically clustered as interneurons. Cells within ±80 µm of the defined *st. pyr* channel were manually classified by a reviewer blind to animal ID and sham/injured condition based on firing rate, spike shape, and autocorrelogram.

Automated k-means consensus clustering was performed to further validate manual clustering. Firing rates, spike width (width of mean action potential waveform at half-max spike height), and the first moment of the autocorrelogram (mean of autocorrelogram values at lags from 0-50 ms) were normalized from 0-1 for each cell within ±80 µm of the defined *st. pyr* channel. K-means clustering (with 2 groups) was performed on these normalized features 1000 times, and the consensus of all 1000 permutations was used to identify cells as pyramidal cells or interneurons. Classification of cells that were manually identified as pyramidal cells or interneurons (excluding unclassified cells) were compared to automated clustering results and there was >95% agreement between the two methods.

### Bursting

Interspike intervals were calculated by subtracting consecutive spike timestamps from each other and bursts were defined as 2 or more spikes within a period of 9 milliseconds. The burst probability was defined as the probability that a cell fired a burst vs a single action potential: burst probability = number of bursts/(number or bursts + number of single spikes outside of bursts).

### Entrainment Analyses

For all entrainment analyses, spikes were referenced to the LFP from the defined *st. pyr* channel on the shank which the unit was recorded from. Entrainment was assessed by filtering the *st. pyr* LFP into discrete frequency bands using a 2^nd^ order Butterworth filter, finding the instantaneous angle (peak=0°) associated with each spike using the Hilbert transform, then computing the mean vector length (MVL) and mean angle of entrainment using circular statistics (CircStat toolbox in MATLAB; RRID:SCR_016651)^159^. Entrainment strength was visualized across a range of frequency bins (24 bins spanning 1-15 Hz with a width of 1 Hz, and 30 bins spanning 15-180 Hz with a width of 20 Hz) separately for epochs when rats were moving (>10 cm/sec) and still (<10 cm/sec) (**Fig 4B**). Units were excluded if their firing rate was below 0.1 Hz in the environment or if they did not have at least 25 spikes in both moving and still epochs.

Theta and gamma entrainment was assessed by filtering the *st. pyr* LFP from 5-10 Hz or 30-59 Hz respectively and calculating the MVL and mean angle of entrainment. To determine if cells were significantly entrained to theta or gamma oscillations (**Fig 4C**), we used a circular shuffling procedure with 500 permutations. For each permutation, the vector of *st. pyr* theta and gamma phase angles was cut at a random point (minimum 10 second offset from the original data), and the phase angles after the cut point were placed in front of the phase angles before the cut point to create a circularly shuffled distribution of phase angles. This maintains the spiking profile of all units as well as the temporal structure of the data except at the cut point. The MVL was computed for each permutation, and significantly entrained cells were defined as cells that had a MVL greater than 95% of shuffled permutations. Any units not significantly entrained to theta oscillations were excluded from analysis in **Fig 4D-F** and **Fig 5**. Theta matched entrainment analyses (**Fig 5A&C**) were computed using a deterministic method that was run independently on interneurons and pyramidal cells. First, theta amplitudes at the time of each spike were extracted. We then looped through each spike from injured animals and found a corresponding spike from a sham animal that had the closet theta amplitude. These spikes were then separated from the dataset, and the process was repeated until the difference in theta amplitudes between spikes exceeded a threshold of 0.01 mV. Only spikes that were amplitude matched between groups were included, and any cells that did not have at least 25 amplitude matched spikes were excluded from analysis. Theta entrainment was calculated for the remaining cells as described above using only the amplitude matched spikes and compared across groups. For theta amplitude vs MVL correlations, a distribution of theta amplitudes at each spike was created independently for interneurons and pyramidal cells. Distributions were then split into bins containing an equal number of spikes (**Fig 5B&D**), and the MVL was calculated in each bin. Any cells that did not have at least 25 spikes in each bin were excluded from analysis. In order to maximize the number of cells included in the analysis while maintaining enough bins to assess the relationship between theta amplitude and MVL, we used 5 bins for interneurons, but only 4 bins for pyramidal cells due to their overall lower firing rates. The average MVL was calculated across each bin, and a line was fit to the data using linear regression.

### Ripple Detection and Analysis

Raw signals were down sampled to 3000 Hz using an anti-aliasing filter (resample function in MATLAB). Then a continuous wavelet transform was used to construct time-frequency maps of signals using an open-source algorithm (fCWT)^160^. Morlet wavelets with 𝜎 = 12 were used for higher frequency resolution. High frequency events were isolated by thresholding the time-frequency maps using empirical cumulative distribution function of amplitudes. The thresholding value was calculated for 80-250 Hz and 250-500 Hz separately to minimize the effect of the decrease in the amplitude of oscillations with frequency. These isolated high frequency blobs were accepted if they passed several criteria such as their duration (at least 3 cycles based on the frequency of oscillation), or their amplitudes compared to background activity. The hyperparameters of this algorithm were chosen by evaluating the accuracy and precision of this method using synthetic data^82^.

Sharp wave events were detected separately on *st. rad* (note that for some animals, the first day of recording did not include any channels located at *st. rad* because most electrodes were localized to *st. pyr*). First, the down sampled raw signals were filtered between 8 and 40 Hz using a 2^nd^ order Butterworth filter. Then, local minimums of the filtered signal were found using islocalmin function in MATLAB as candidates for the center of sharp waves. Next, islocalmax function in MATLAB was used to find the local maximums of the filtered signal for the starting and stopping point of a sharp wave. A sharp wave was identified if a center (local minimum) was located between to local maximums. Finally, sharp waves were only accepted if their amplitude of the local minimum was larger than 98% of all local minimums. The described method for sharp wave detection was generous by design (it can be stricter by applying a harsher criterion for amplitudes).

Sharp-wave ripples were identified if the detected ripples events (accepted contiguous blobs in time-frequency maps) overlapped with detected sharp wave events. These putative SWRs were then manually curated by an observer blind to animal ID, sham/injured condition, and animal behavior during event. The percentage of accepted SWR events that occurred while the animals were not moving was calculated (>95% in all conditions) and SWR event frequency was normalized to the amount of time animals were not moving to compare SWR event frequency across conditions while controlling for any potential differences is the amount of time animals spent moving. Frequency amplitude spectra were computed for each event by averaging amplitudes at each frequency bin over times coinciding with the blob edges (see white lines in **Fig 6A**), then averaged across conditions. Event durations and peak amplitudes (the maximum value in the time-frequency map) were then computed to create the cumulative distributions shown in **Fig 6E**.

### Morris Water Maze

The MWM apparatus was a circular pool (2 m in diameter, 50 cm tall, filled to a depth of 25 cm with 18° C water) containing a hidden platform (11.5 cm x 11.5 cm x 24 cm tall) just below the water’s surface. Both the interior of the pool and the platform were painted black making the platform invisible from the surface. External visual cues could be easily seen from the pool for navigation purposes. Rats received 10 trials of training in the MWM ∼20 hrs before TBI/sham surgery followed by another 10 trials ∼2 hrs before surgery. For each trial, the rat was placed in one of 4 randomly selected starting locations located 90° apart and given 1 min to freely swim until they found the platform. If the rat was unable to find the platform after 1 min, they were placed there for 15-30 sec. Rats were tested ∼42 hrs after FPI or sham surgery to assess memory of the platform location. During testing, the platform was removed, and rats had 2 testing trials during which they freely swam for 1 minute. A memory score^86^ was calculated based on the amount of time the rat spent in 5 distinct zones, with zones closer to the platform’s previous location being weighted higher. All testing trials were video recorded from a camera placed above the pool, and the rat’s position was obtained using automated video tracking (Ethovision XT, Noduls; RRID: SCR_000441). Video recordings were obtained during training from a subset of animals (7/9 sham and 10/14 injured) and the latency to reach the platform was determined. If a rat was unable to find the platform in 1 min, as described above, the latency for that trial was set to 60 sec.

### Histological Examinations

To ensure the nature and severity of injury were consistent with prior characterizations of the model, a separate set of animals not implanted with electrodes underwent FPI (n=2; 1.8 atm) or sham surgery (n=1) for histopathological analysis. All animals were survived for 48 hours. At the study endpoint, ketamine / xylazine / acepromazine (75/15/2 mg/kg, i.p.) was administered and following adequate anesthesia, animals were transcardially perfused with chilled 1X phosphate buffered saline (PBS) followed immediately with chilled 10% neutral buffered formalin. Brains were extracted and post-fixed for 24 hours at 4°C before being blocked in the coronal plane at 2 mm intervals and processed to paraffin using standard techniques. 8 µm thick whole brain coronal sections were obtained at the level of the anterior hippocampus, posterior hippocampus and posterior angular bundle using a rotary microtome.

All regions were stained using hematoxylin and eosin (H&E). In addition, to determine the presence and distribution of axonal pathology, immunohistochemical (IHC) techniques were performed as previously described^92,161^. Briefly, following deparaffinization and rehydration of tissue sections, endogenous peroxidase activity tissue was quenched using 3% aqueous hydrogen peroxide. Antigen retrieval was performed using a pressure cooker with sections immersed in Tris EDTA buffer (pH 8.0). Subsequent blocking was performed for 30 minutes in 1% normal horse serum (Vector Labs, Burlingame, CA, USA) in Optimax buffer (BioGenex, San Ramon, CA, USA). An antibody reactive for the N-terminal amino acids 66-81 of the amyloid precursor protein (APP) (Clone 22C11: Millipore, Billerica, MA) was applied at 1:40K and incubated overnight at 4° C. After rinsing, sections were incubated with the appropriate biotinylated secondary antibody for 30 minutes followed by avidin-biotin complex and visualization achieved using DAB (all reagents: Vector Labs, Burlingame, CA, USA). Sections were rinsed and dehydrated in graded alcohols and cleared in xylenes before being coverslipped. Positive control tissue for APP IHC included sections of contused rat brain tissue with previously established axonal pathology. Omission of the primary antibody was performed on positive control tissue to control for non-specific binding.

Fluoro-Jade C staining was performed to assess for neuronal degeneration using the Biosensis Fluoro-Jade C staining Reagent Kit per the manufacturer’s instructions (Biosensis, Thebarton, Australia). Briefly, following dewaxing and rehydration to water as above, tissue was immersed in potassium permanganate solution for 25 minutes at room temperature. After being rinsed in gently flowing dH_2_O, tissue was then incubated in Fluoro-Jade C solution at room temperature for 30 minutes. After rinsing, sections were dried at 37° C in an oven for 90 minutes before being immersed in xylenes and coverslipped.

### Analysis of Histological Findings

All sections were reviewed for the presence of hemorrhage. The presence or absence of axonal pathology was determined in all sections including in the fimbria fornix, angular bundle, hippocampus, corpus callosum and thalamus. In addition, cell death was assessed by examination for Fluoro-Jade C positive neurons in all sections, including all hippocampal subfields, entorhinal cortex and thalamus. Two independent observers reviewed all sections for the presence or absence of axonal pathology and cell degeneration with excellent interrater reliability (Cohen’s Kappa 0.91). Notably, histological observations were performed at 48 hours to identify pathologies which are readily visualized at this timepoint including acute axonal and neuronal degeneration. However, due to the complex temporal evolution of post-traumatic pathologies, observations at 48 hrs may differ from the pathologies at 7 days post-injury, when electrophysiological recordings were performed.

## AUTHOR CONTRIBUTIONS

Conceptualization: CDA, CC, KGG, EM, JAW

Formal Analysis: CDA, JDA, CC, KGG, EM

Investigation: CDA, KGG

Resources: VEJ, JAW

Writing – Original Draft Preparation: CDA

Writing – Review & Editing: CDA, JDA, VEJ, EM, AVU, JAW

Visualization: CDA, JDA, EM

Supervision: VEJ, JAW

Funding Acquisition: JAW

## REFERENCES

Draper, K. & Ponsford, J. Cognitive Functioning Ten Years Following Traumatic Brain Injury and Rehabilitation. Neuropsychology 22, 618–625, doi:10.1037/0894-4105.22.5.618 (2008).

Monti, J. M. et al. History of mild traumatic brain injury is associated with deficits in relational memory, reduced hippocampal volume, and less neural activity later in life. Front. Aging Neurosci. 5, 41, doi:10.3389/fnagi.2013.00041 (2013).

Vakil, E., Greenstein, Y., Weiss, I. & Shtein, S. The Effects of Moderate-to-Severe Traumatic Brain Injury on Episodic Memory: a Meta-Analysis. Neuropsychol Rev 29, 270–287, doi:10.1007/s11065-019-09413-8 (2019).

Bigler, E. D. et al. Traumatic Brain Injury and Memory: The Role of Hippocampal Atrophy. Neuropsychology 10, 333–342, doi:10.1037/0894-4105.10.3.333 (1996).

Eakin, K. & Miller, J. P. Mild Traumatic Brain Injury Is Associated with Impaired Hippocampal Spatiotemporal Representation in the Absence of Histological Changes. Journal of Neurotrauma 29, 1180–1187, doi:10.1089/neu.2011.2192 (2012).

Lyeth, B. G. et al. Prolonged memory impairment in the absence of hippocampal cell death following traumatic brain injury in the rat. Brain Research 526, 249–258, doi:10.1016/0006-8993(90)91229-a (1990).

Tate, D. F. & Bigler, E. D. Fornix and Hippocampal Atrophy in Traumatic Brain Injury. Learn. Mem. 7, 442–446, doi:10.1101/lm.33000 (2000).

Adnan, A. et al. Moderate–severe traumatic brain injury causes delayed loss of white matter integrity: Evidence of fornix deterioration in the chronic stage of injury. Brain Inj. 27, 1415–1422, doi:10.3109/02699052.2013.823659 (2013).

Blumbergs, P. C. et al. Stalning af amyloid percursor protein to study axonal damage in mild head Injury. Lancet 344, 1055–1056, doi:10.1016/s0140-6736(94)91712-4 (1994).

Kinnunen, K. M. et al. White matter damage and cognitive impairment after traumatic brain injury. Brain 134, 449–463, doi:10.1093/brain/awq347 (2011).

Palacios, E. M. et al. Diffusion tensor imaging differences relate to memory deficits in diffuse traumatic brain injury. BMC Neurol. 11, 24, doi:10.1186/1471-2377-11-24 (2011).

Tomaiuolo, F. et al. Gross morphology and morphometric sequelae in the hippocampus, fornix, and corpus callosum of patients with severe non-missile traumatic brain injury without macroscopically detectable lesions: a T1 weighted MRI study. *J. Neurol., Neurosurg*. Psychiatry 75, 1314, doi:10.1136/jnnp.2003.017046 (2004).

Faden, A. I., Demediuk, P., Panter, S. S. & Vink, R. The role of excitatory amino acids and NMDA receptors in traumatic brain injury. Science 244, 798–800, doi:10.1126/science.2567056 (1989).

Guerriero, R. M., Giza, C. C. & Rotenberg, A. Glutamate and GABA Imbalance Following Traumatic Brain Injury. Curr. Neurol. Neurosci. Rep. 15, 27, doi:10.1007/s11910-015-0545-1 (2015).

McGuire, J. L., Ngwenya, L. B. & McCullumsmith, R. E. Neurotransmitter changes after traumatic brain injury: an update for new treatment strategies. Mol. Psychiatry 24, 995–1012, doi:10.1038/s41380-018-0239-6 (2019).

Aungst, S. L., Kabadi, S. V., Thompson, S. M., Stoica, B. A. & Faden, A. I. Repeated Mild Traumatic Brain Injury Causes Chronic Neuroinflammation, Changes in Hippocampal Synaptic Plasticity, and Associated Cognitive Deficits. J. Cereb. Blood Flow Metab. 34, 1223–1232, doi:10.1038/jcbfm.2014.75 (2014).

Muccigrosso, M. M. et al. Cognitive deficits develop 1month after diffuse brain injury and are exaggerated by microglia-associated reactivity to peripheral immune challenge. *Brain, Behav.*, Immun. 54, 95–109, doi:10.1016/j.bbi.2016.01.009 (2016).

Shetty, A. K., Mishra, V., Kodali, M. & Hattiangady, B. Blood brain barrier dysfunction and delayed neurological deficits in mild traumatic brain injury induced by blast shock waves. Front. Cell. Neurosci. 8, 232, doi:10.3389/fncel.2014.00232 (2014).

Sulhan, S., Lyon, K. A., Shapiro, L. A. & Huang, J. H. Neuroinflammation and blood–brain barrier disruption following traumatic brain injury: Pathophysiology and potential therapeutic targets. J. Neurosci. Res. 98, 19–28, doi:10.1002/jnr.24331 (2020).

Lai, J.-q., Shi, Y.-C., Lin, S. & Chen, X.-R. Metabolic disorders on cognitive dysfunction after traumatic brain injury. Trends Endocrinol. Metab. 33, 451–462, doi:10.1016/j.tem.2022.04.003 (2022).

Li, J. et al. Exploring Temporospatial Changes in Glucose Metabolic Disorder, Learning, and Memory Dysfunction in a Rat Model of Diffuse Axonal Injury. Journal of Neurotrauma 29, 2635–2646, doi:10.1089/neu.2012.2411 (2012).

Weil, Z. M., Gaier, K. R. & Karelina, K. Injury timing alters metabolic, inflammatory and functional outcomes following repeated mild traumatic brain injury. Neurobiology of Disease 70, 108–116, doi:10.1016/j.nbd.2014.06.016 (2014).

Yi, L. et al. Serum Metabolic Profiling Reveals Altered Metabolic Pathways in Patients with Post-traumatic Cognitive Impairments. Sci. Rep. 6, 21320, doi:10.1038/srep21320 (2016).

D’Ambrosio, R., Maris, D. O., Grady, M. S., Winn, H. R. & Janigro, D. Selective loss of hippocampal long-term potentiation, but not depression, following fluid percussion injury. Brain Research 786, 64–79, doi:10.1016/s0006-8993(97)01412-1 (1998).

Folweiler, K. A., Samuel, S., Metheny, H. E. & Cohen, A. S. Diminished Dentate Gyrus Filtering of Cortical Input Leads to Enhanced Area Ca3 Excitability after Mild Traumatic Brain Injury. Journal of Neurotrauma 35, 1304–1317, doi:10.1089/neu.2017.5350 (2018).

Katayama, Y., Becker, D. P., Tamura, T. & Hovda, D. A. Massive increases in extracellular potassium and the indiscriminate release of glutamate following concussive brain injury. Journal of Neurosurgery 73, 889–900, doi:10.3171/jns.1990.73.6.0889 (1990).

Lei, Z., Deng, P., Li, J. & Xu, Z. C. Alterations of A-Type Potassium Channels in Hippocampal Neurons after Traumatic Brain Injury. Journal of Neurotrauma 29, 235–245, doi:10.1089/neu.2010.1537 (2012).

Miyazaki, S. et al. Enduring suppression of hippocampal long-term potentiation following traumatic brain injury in rat. Brain Research 585, 335–339, doi:10.1016/0006-8993(92)91232-4 (1992).

Reeves, T. M., Lyeth, B. G. & Povlishock, J. T. Long-term potentiation deficits and excitability changes following traumatic brain injury. Exp Brain Res 106, 248–256, doi:10.1007/bf00241120 (1995).

Sanders, M. J., Sick, T. J., Perez-Pinzon, M. A., Dietrich, W. D. & Green, E. J. Chronic failure in the maintenance of long-term potentiation following fluid percussion injury in the rat. Brain Research 861, 69–76, doi:10.1016/s0006-8993(00)01986-7 (2000).

Santhakumar, V., Ratzliff, A. D. H., Jeng, J., Toth, Z. & Soltesz, I. Long-term hyperexcitability in the hippocampus after experimental head trauma. Ann Neurol 50, 708–717, doi:10.1002/ana.1230 (2001).

Schwarzbach, E., Bonislawski, D. P., Xiong, G. & Cohen, A. S. Mechanisms underlying the inability to induce area CA1 LTP in the mouse after traumatic brain injury. Hippocampus 16, 541–550, doi:10.1002/hipo.20183 (2006).

Witgen, B. M. et al. Regional hippocampal alteration associated with cognitive deficit following experimental brain injury: A systems, network and cellular evaluation. Neuroscience 133, 1–15, doi:10.1016/j.neuroscience.2005.01.052 (2005).

Wolf, J. A. et al. Concussion Induces Hippocampal Circuitry Disruption in Swine. Journal of Neurotrauma 34, 2303–2314, doi:10.1089/neu.2016.4848 (2017).

Huang, Y., Brandon, M. P., Griffin, A. L., Hasselmo, M. E. & Eden, U. T. Decoding Movement Trajectories Through a T-Maze Using Point Process Filters Applied to Place Field Data from Rat Hippocampal Region CA1. Neural Comput. 21, 3305–3334, doi:10.1162/neco.2009.10-08-893 (2009).

Lisman, J. & Redish, A. D. Prediction, sequences and the hippocampus. Philos. Trans. R. Soc. B: Biol. Sci. 364, 1193–1201, doi:10.1098/rstb.2008.0316 (2009).

Pastalkova, E., Itskov, V., Amarasingham, A. & Buzsáki, G. r. Internally Generated Cell Assembly Sequences in the Rat Hippocampus. Science 321, 1322–1327, doi:10.1126/science.1159775 (2008).

Wikenheiser, A. M. & Redish, A. D. Hippocampal theta sequences reflect current goals. Nat Neurosci 18, 289–294, doi:10.1038/nn.3909 (2015).

Xu, H., Baracskay, P., O’Neill, J. & Csicsvari, J. Assembly Responses of Hippocampal CA1 Place Cells Predict Learned Behavior in Goal-Directed Spatial Tasks on the Radial Eight-Arm Maze. Neuron 101, 119–132.e114, doi:10.1016/j.neuron.2018.11.015 (2019).

Biswas, C., Marković, D. & Giza, C. C. Alterations in Mesoscopic Oscillations affecting Episodic Memory following Developmental Traumatic Brain Injury. Experimental Neurology 300, 259–273, doi:10.1016/j.expneurol.2017.10.021 (2018).

Broussard, J. I. et al. Mild Traumatic Brain Injury Decreases Spatial Information Content and Reduces Place Field Stability of Hippocampal CA1 Neurons. Journal of Neurotrauma 37, 227–235, doi:10.1089/neu.2019.6766 (2020).

Fedor, M., Berman, R. F., Muizelaar, J. P. & Lyeth, B. G. Hippocampal Theta Dysfunction after Lateral Fluid Percussion Injury. Journal of Neurotrauma 27, 1605–1615, doi:10.1089/neu.2010.1370 (2010).

Koch, P. F. et al. Traumatic Brain Injury Preserves Firing Rates but Disrupts Laminar Oscillatory Coupling and Neuronal Entrainment in Hippocampal CA1. Eneuro 7, ENEURO.0495-0419.2020, doi:10.1523/eneuro.0495-19.2020 (2020).

Munyon, C., Eakin, K. C., Sweet, J. A. & Miller, J. P. Decreased bursting and novel object-specific cell firing in the hippocampus after mild traumatic brain injury. Brain Research 1582, 220–226, doi:10.1016/j.brainres.2014.07.036 (2014).

Paterno, R., Metheny, H., Xiong, G., Elkind, J. & Cohen, A. S. Mild Traumatic Brain Injury Decreases Broadband Power in Area CA1. Journal of Neurotrauma 33, 1645–1649, doi:10.1089/neu.2015.4107 (2016).

Noble, B. et al. Mild Traumatic Brain Injury Impairs Coupling of CA1 Neuronal Activity to Theta Oscillations. Neurotrauma Rep. 5, 1079–1088, doi:10.1089/neur.2024.0119 (2024).

Fernández-Ruiz, A., et al. Entorhinal-CA3 Dual-Input Control of Spike Timing in the Hippocampus by Theta-Gamma Coupling. Neuron 93, 1213–1226.e1215, doi:10.1016/j.neuron.2017.02.017 (2017).

Hasselmo, M. E., Bodeln, C. & Wyble, B. P. A Proposed Function for Hippocampal Theta Rhythm: Separate Phases of Encoding and Retrieval Enhance Reversal of Prior Learning. Neural Comput. 14, 793–817, doi:10.1162/089976602317318965 (2002).

Feldman, Daniel E. The Spike-Timing Dependence of Plasticity. Neuron 75, 556–571, doi:10.1016/j.neuron.2012.08.001 (2012).

Sirota, A. et al. Entrainment of Neocortical Neurons and Gamma Oscillations by the Hippocampal Theta Rhythm. Neuron 60, 683–697, doi:10.1016/j.neuron.2008.09.014 (2008).

Buzsáki, G. Theta Oscillations in the Hippocampus. Neuron 33, 325–340, doi:10.1016/s0896-6273(02)00586-x (2002).

Vanderwolf, C. H. Hippocampal electrical activity and voluntary movement in the rat. Electroencephalogr. Clin. Neurophysiol. 26, 407–418, doi:10.1016/0013-4694(69)90092-3 (1969).

Solomon, E. A. et al. Widespread theta synchrony and high-frequency desynchronization underlies enhanced cognition. Nature Communications 8, 1704, doi:10.1038/s41467-017-01763-2 (2017).

Summerfield, C. & Mangels, J. A. Coherent theta-band EEG activity predicts item-context binding during encoding. NeuroImage 24, 692–703, doi:10.1016/j.neuroimage.2004.09.012 (2005).

Csicsvari, J., Jamieson, B., Wise, K. D. & Buzsáki, G. Mechanisms of Gamma Oscillations in the Hippocampus of the Behaving Rat. Neuron 37, 311–322, doi:10.1016/s0896-6273(02)01169-8 (2003).

Bragin, A. et al. Gamma (40-100 Hz) oscillation in the hippocampus of the behaving rat. The Journal of Neuroscience 15, 47–60, doi:10.1523/jneurosci.15-01-00047.1995 (1995).

Buzsáki, G., S, L. L.-W. & Vanderwolf, C. H. Cellular bases of hippocampal EEG in the behaving rat. Brain Res. Rev. 6, 139–171, doi:10.1016/0165-0173(83)90037-1 (1983).

Canolty, R. T. et al. High Gamma Power Is Phase-Locked to Theta Oscillations in Human Neocortex. Science 313, 1626–1628, doi:10.1126/science.1128115 (2006).

Colgin, L. L. Theta–gamma coupling in the entorhinal–hippocampal system. Current Opinion in Neurobiology 31, 45–50, doi:10.1016/j.conb.2014.08.001 (2015).

Jensen, O. & Colgin, L. L. Cross-frequency coupling between neuronal oscillations. Trends Cogn Sci 11, 267–269, doi:10.1016/j.tics.2007.05.003 (2007).

Lega, B., Burke, J., Jacobs, J. & Kahana, M. J. Slow-Theta-to-Gamma Phase–Amplitude Coupling in Human Hippocampus Supports the Formation of New Episodic Memories. Cereb. Cortex 26, 268–278, doi:10.1093/cercor/bhu232 (2016).

Axmacher, N. et al. Cross-frequency coupling supports multi-item working memory in the human hippocampus. Proceedings of the National Academy of Sciences 107, 3228–3233, doi:10.1073/pnas.0911531107 (2010).

Daume, J. et al. Control of working memory by phase–amplitude coupling of human hippocampal neurons. Nature, 1–9, doi:10.1038/s41586-024-07309-z (2024).

Buzsáki, G. Hippocampal sharp waves: Their origin and significance. Brain Research 398, 242–252, doi:10.1016/0006-8993(86)91483-6 (1986).

Buzsáki, G. Hippocampal sharp wave-ripple: A cognitive biomarker for episodic memory and planning. Hippocampus 25, 1073–1188, doi:10.1002/hipo.22488 (2015).

Carr, M. F., Jadhav, S. P. & Frank, L. M. Hippocampal replay in the awake state: a potential substrate for memory consolidation and retrieval. Nat Neurosci 14, 147–153, doi:10.1038/nn.2732 (2011).

Davidson, T. J., Kloosterman, F. & Wilson, M. A. Hippocampal Replay of Extended Experience. Neuron 63, 497–507, doi:10.1016/j.neuron.2009.07.027 (2009).

Wilson, M. A. & McNaughton, B. L. Reactivation of Hippocampal Ensemble Memories During Sleep. Science 265, 676–679, doi:10.1126/science.8036517 (1994).

Donoghue, T. et al. Parameterizing neural power spectra into periodic and aperiodic components. Nat Neurosci 23, 1655–1665, doi:10.1038/s41593-020-00744-x (2020).

Wolf, J. A. & Koch, P. F. Disruption of Network Synchrony and Cognitive Dysfunction After Traumatic Brain Injury. Frontiers in Systems Neuroscience 10, 43, doi:10.3389/fnsys.2016.00043 (2016).

Tort, A. B. L. et al. Dynamic cross-frequency couplings of local field potential oscillations in rat striatum and hippocampus during performance of a T-maze task. Proceedings of the National Academy of Sciences 105, 20517–20522, doi:10.1073/pnas.0810524105 (2008).

Csicsvari, J., Hirase, H., Czurko, A. & Buzsáki, G. Reliability and State Dependence of Pyramidal Cell–Interneuron Synapses in the Hippocampus an Ensemble Approach in the Behaving Rat. Neuron 21, 179–189, doi:10.1016/s0896-6273(00)80525-5 (1998).

Tummala, S. R., Hemphill, M. A., Nam, A. & Meaney, D. F. Concussion increases CA1 activity during prolonged inactivity in a familiar environment. Experimental Neurology 334, 113435, doi:10.1016/j.expneurol.2020.113435 (2020).

Lasztóczi, B. & Klausberger, T. Layer-Specific GABAergic Control of Distinct Gamma Oscillations in the CA1 Hippocampus. Neuron 81, 1126–1139, doi:10.1016/j.neuron.2014.01.021 (2014).

Royer, S. et al. Control of timing, rate and bursts of hippocampal place cells by dendritic and somatic inhibition. Nat Neurosci 15, 769–775, doi:10.1038/nn.3077 (2012).

Vida, I., Bartos, M. & Jonas, P. Shunting Inhibition Improves Robustness of Gamma Oscillations in Hippocampal Interneuron Networks by Homogenizing Firing Rates. Neuron 49, 107–117, doi:10.1016/j.neuron.2005.11.036 (2006).

Dragoi, G. & Buzsáki, G. Temporal Encoding of Place Sequences by Hippocampal Cell Assemblies. Neuron 50, 145–157, doi:10.1016/j.neuron.2006.02.023 (2006).

Foster, D. J. & Wilson, M. A. Hippocampal theta sequences. Hippocampus 17, 1093–1099, doi:10.1002/hipo.20345 (2007).

Gupta, A. S., Meer, M. A. A. v. d., Touretzky, D. S. & Redish, A. D. Segmentation of spatial experience by hippocampal theta sequences. Nat Neurosci 15, 1032–1039, doi:10.1038/nn.3138 (2012).

O’Keefe, J. & Recce, M. L. Phase relationship between hippocampal place units and the EEG theta rhythm. Hippocampus 3, 317–330, doi:10.1002/hipo.450030307 (1993).

Skaggs, W. E., McNaughton, B. L., Wilson, M. A. & Barnes, C. A. Theta phase precession in hippocampal neuronal populations and the compression of temporal sequences. Hippocampus 6, 149–172, doi:10.1002/(sici)1098-1063(1996)6:2<149::aid-hipo6>3.0.co;2-k (1996).

Mirzakhalili, E., Adam, C. D., Ulyanova, A. V., Johnson, V. E. & Wolf, J. A. Automatic High-Frequency Oscillations Detection Using Time-Frequency Analysis. 2023 11th Int Ieee Embs Conf Neural Eng Ner 00, 1–6, doi:10.1109/ner52421.2023.10123882 (2023).

Bramlett, H. M. & Dietrich, D. W. Quantitative structural changes in white and gray matter 1 year following traumatic brain injury in rats. Acta Neuropathol. 103, 607–614, doi:10.1007/s00401-001-0510-8 (2002).

McIntosh, T. K. et al. Traumatic brain injury in the rat: Characterization of a lateral fluid-percussion model. Neuroscience 28, 233–244, doi:10.1016/0306-4522(89)90247-9 (1989).

Saatman, K. E., Graham, D. I. & McIntosh, T. K. The Neuronal Cytoskeleton Is at Risk After Mild and Moderate Brain Injury. Journal of Neurotrauma 15, 1047–1058, doi:10.1089/neu.1998.15.1047 (1998).

Smith, D. H., Okiyama, K., Thomas, M. J., Claussen, B. & McIntosh, T. K. Evaluation of Memory Dysfunction Following Experimental Brain Injury Using the Morris Water Maze. Journal of Neurotrauma 8, 259–269, doi:10.1089/neu.1991.8.259 (1991).

Thompson, H. J. et al. Lateral Fluid Percussion Brain Injury: A 15-Year Review and Evaluation. Journal of Neurotrauma 22, 42–75, doi:10.1089/neu.2005.22.42 (2005).

Ndode-Ekane, X. E. et al. Harmonization of lateral fluid-percussion injury model production and post-injury monitoring in a preclinical multicenter biomarker discovery study on post-traumatic epileptogenesis. Epilepsy Res. 151, 7–16, doi:10.1016/j.eplepsyres.2019.01.006 (2019).

Vink, R., Mullins, P. G. M., Temple, M. D., Bao, W. & Faden, A. I. Small Shifts in Craniotomy Position in the Lateral Fluid Percussion Injury Model Are Associated with Differential Lesion Development. Journal of Neurotrauma 18, 839–847, doi:10.1089/089771501316919201 (2001).

Carlson, S. W., Henchir, J. & Dixon, C. E. Lateral Fluid Percussion Injury Impairs Hippocampal Synaptic Soluble N-Ethylmaleimide Sensitive Factor Attachment Protein Receptor Complex Formation. Front. Neurol. 8, 532, doi:10.3389/fneur.2017.00532 (2017).

Gentleman, S. M., Nash, M. J., Sweeting, C. J., Graham, D. I. & Roberts, G. W. β-Amyloid precursor protein (βAPP) as a marker for axonal injury after head injury. Neurosci. Lett. 160, 139–144, doi:10.1016/0304-3940(93)90398-5 (1993).

Johnson, V. E. et al. Inflammation and white matter degeneration persist for years after a single traumatic brain injury. Brain 136, 28–42, doi:10.1093/brain/aws322 (2013).

Johnson, V. E., Stewart, W. & Smith, D. H. Axonal pathology in traumatic brain injury. Experimental Neurology 246, 35–43, doi:10.1016/j.expneurol.2012.01.013 (2013).

Johnson, V. E. et al. SNTF immunostaining reveals previously undetected axonal pathology in traumatic brain injury. Acta Neuropathol. 131, 115–135, doi:10.1007/s00401-015-1506-0 (2016).

Freund, T. F. & Buzsáki, G. Interneurons of the hippocampus. Hippocampus 6, 347–470, doi:10.1002/(sici)1098-1063(1996)6:4<347::aid-hipo1>3.0.co;2-i (1996).

Pelkey, K. A. et al. Hippocampal GABAergic Inhibitory Interneurons. Physiol Rev 97, 1619–1747, doi:10.1152/physrev.00007.2017 (2017).

Schlingloff, D., Káli, S., Freund, T. F., Hájos, N. & Gulyás, A. I. Mechanisms of Sharp Wave Initiation and Ripple Generation. The Journal of Neuroscience 34, 11385–11398, doi:10.1523/jneurosci.0867-14.2014 (2014).

Stark, E. et al. Pyramidal Cell-Interneuron Interactions Underlie Hippocampal Ripple Oscillations. Neuron 83, 467–480, doi:10.1016/j.neuron.2014.06.023 (2014).

Ylinen, A. et al. Sharp wave-associated high-frequency oscillation (200 Hz) in the intact hippocampus: network and intracellular mechanisms. Journal of Neuroscience 15, 30–46, doi:10.1523/jneurosci.15-01-00030.1995 (1995).

Almeida-Suhett, C. P. et al. GABAergic interneuronal loss and reduced inhibitory synaptic transmission in the hippocampal CA1 region after mild traumatic brain injury. Experimental Neurology 273, 11–23, doi:10.1016/j.expneurol.2015.07.028 (2015).

Frankowski, J. C., Kim, Y. J. & Hunt, R. F. Selective vulnerability of hippocampal interneurons to graded traumatic brain injury. Neurobiology of Disease 129, 208–216, doi:10.1016/j.nbd.2018.07.022 (2018).

Ulyanova, A. V. et al. Hippocampal interneuronal dysfunction and hyperexcitability in a porcine model of concussion. *Commun*. Biol. 6, 1136, doi:10.1038/s42003-023-05491-w (2023).

Cortez, S. C., McIntosh, T. K. & Noble, L. J. Experimental fluid percussion brain injury: vascular disruption and neuronal and glial alterations. Brain Research 482, 271–282, doi:10.1016/0006-8993(89)91190-6 (1989).

Hicks, R. R., Smith, D. H., Lowenstein, D. H., Marie, R. S. & McIntosh, T. K. Mild Experimental Brain Injury in the Rat Induces Cognitive Deficits Associated with Regional Neuronal Loss in the Hippocampus. Journal of Neurotrauma 10, 405–414, doi:10.1089/neu.1993.10.405 (1993).

Maxwell, W. L. et al. There Is Differential Loss of Pyramidal Cells from the Human Hippocampus with Survival after Blunt Head Injury. J. Neuropathol. Exp. Neurol. 62, 272–279, doi:10.1093/jnen/62.3.272 (2003).

Swartz, B. E. et al. Hippocampal Cell Loss in Posttraumatic Human Epilepsy. Epilepsia 47, 1373–1382, doi:10.1111/j.1528-1167.2006.00602.x (2006).

Christidi, F. et al. Diffusion Tensor Imaging of the Perforant Pathway Zone and Its Relation to Memory Function in Patients with Severe Traumatic Brain Injury. Journal of Neurotrauma 28, 711–725, doi:10.1089/neu.2010.1644 (2011).

Colley, B. S., Phillips, L. L. & Reeves, T. M. The effects of cyclosporin-A on axonal conduction deficits following traumatic brain injury in adult rats. Experimental Neurology 224, 241–251, doi:10.1016/j.expneurol.2010.03.026 (2010).

Reeves, T. M., Phillips, L. L. & Povlishock, J. T. Myelinated and unmyelinated axons of the corpus callosum differ in vulnerability and functional recovery following traumatic brain injury. Experimental Neurology 196, 126–137, doi:10.1016/j.expneurol.2005.07.014 (2005).

Deng, P. & Xu, Z. C. Contribution of Ih to Neuronal Damage in the Hippocampus after Traumatic Brain Injury in Rats. Journal of Neurotrauma 28, 1173–1183, doi:10.1089/neu.2010.1683 (2011).

Karimi, S. A., Hosseinmardi, N., Sayyah, M., Hajisoltani, R. & Janahmadi, M. Enhancement of intrinsic neuronal excitability-mediated by a reduction in hyperpolarization-activated cation current (Ih) in hippocampal CA1 neurons in a rat model of traumatic brain injury. Hippocampus 31, 156–169, doi:10.1002/hipo.23270 (2021).

Sizemore, G. et al. Temporal Lobe Epilepsy, Stroke, and Traumatic Brain Injury: Mechanisms of Hyperpolarized, Depolarized, and Flow-Through Ion Channels Utilized as Tri-Coordinate Biomarkers of Electrophysiologic Dysfunction. OBM Neurobiol. 2, 1–1, doi:10.21926/obm.neurobiol.1802009 (2018).

Leonard, J. R., Grady, M. S., Lee, M. E., Paz, J. C. & Westrum, L. E. Fluid Percussion Injury Causes Disruption of the Septohippocampal Pathway in the Rat. Experimental Neurology 143, 177–187, doi:10.1006/exnr.1996.6366 (1997).

Borhegyi, Z., Varga, V., Szilágyi, N., Fabo, D. & Freund, T. F. Phase Segregation of Medial Septal GABAergic Neurons during Hippocampal Theta Activity. The Journal of Neuroscience 24, 8470–8479, doi:10.1523/jneurosci.1413-04.2004 (2004).

Hangya, B., Borhegyi, Z., Szilagyi, N., Freund, T. F. & Varga, V. GABAergic Neurons of the Medial Septum Lead the Hippocampal Network during Theta Activity. Journal of Neuroscience 29, 8094–8102, doi:10.1523/jneurosci.5665-08.2009 (2009).

Varga, V. et al. The presence of pacemaker HCN channels identifies theta rhythmic GABAergic neurons in the medial septum. J. Physiol. 586, 3893–3915, doi:10.1113/jphysiol.2008.155242 (2008).

Freund, T. F. & Antal, M. GABA-containing neurons in the septum control inhibitory interneurons in the hippocampus. Nature 336, 170–173, doi:10.1038/336170a0 (1988).

Arciniegas, D. B. The cholinergic hypothesis of cognitive impairment caused by traumatic brain injury. Curr Psychiat Rep 5, 391–399, doi:10.1007/s11920-003-0074-5 (2003).

Shin, S. S. & Dixon, C. E. Alterations in Cholinergic Pathways and Therapeutic Strategies Targeting Cholinergic System after Traumatic Brain Injury. Journal of Neurotrauma 32, 1429–1440, doi:10.1089/neu.2014.3445 (2015).

Kramis, R., Vanderwolf, C. H. & Bland, B. H. Two types of hippocampal rhythmical slow activity in both the rabbit and the rat: Relations to behavior and effects of atropine, diethyl ether, urethane, and pentobarbital. Experimental Neurology 49, 58–85, doi:10.1016/0014-4886(75)90195-8 (1975).

Vandecasteele, M. et al. Optogenetic activation of septal cholinergic neurons suppresses sharp wave ripples and enhances theta oscillations in the hippocampus. Proceedings of the National Academy of Sciences 111, 13535–13540, doi:10.1073/pnas.1411233111 (2014).

Vanderwolf, C. H. Neocortical and hippocampal activation in relation to behavior: Effects of atropine, eserine, phenothiazines, and amphetamine. J. Comp. Physiol. Psychol. 88, 300–323, doi:10.1037/h0076211 (1975).

Almeida, L. d., Idiart, M. & Lisman, J. E. A Second Function of Gamma Frequency Oscillations: An E%-Max Winner-Take-All Mechanism Selects Which Cells Fire. The Journal of Neuroscience 29, 7497–7503, doi:10.1523/jneurosci.6044-08.2009 (2009).

Dragoi, G. Cell assemblies, sequences and temporal coding in the hippocampus. Current Opinion in Neurobiology 64, 111–118, doi:10.1016/j.conb.2020.03.003 (2020).

Senior, T. J., Huxter, J. R., Allen, K., O’Neill, J. & Csicsvari, J. Gamma Oscillatory Firing Reveals Distinct Populations of Pyramidal Cells in the CA1 Region of the Hippocampus. The Journal of Neuroscience 28, 2274–2286, doi:10.1523/jneurosci.4669-07.2008 (2008).

Colgin, L. L. et al. Frequency of gamma oscillations routes flow of information in the hippocampus. Nature 462, 353–357, doi:10.1038/nature08573 (2009).

Jezek, K., Henriksen, E. J., Treves, A., Moser, E. I. & Moser, M.-B. Theta-paced flickering between place-cell maps in the hippocampus. Nature 478, 246–249, doi:10.1038/nature10439 (2011).

Hamm, R. J., Lyeth, B. G., Jenkins, L. W., O’Dell, D. M. & Pike, B. R. Selective cognitive impairment following traumatic brain injury in rats. Behav. Brain Res. 59, 169–173, doi:10.1016/0166-4328(93)90164-l (1993).

Broussard, J. I. et al. Optogenetic Stimulation of CA1 Pyramidal Neurons at Theta Enhances Recognition Memory in Brain Injured Animals. Journal of Neurotrauma 40, 2442–2448, doi:10.1089/neu.2023.0078 (2023).

Lee, D. J. et al. Medial Septal Nucleus Theta Frequency Deep Brain Stimulation Improves Spatial Working Memory after Traumatic Brain Injury. Journal of Neurotrauma 30, 131–139, doi:10.1089/neu.2012.2646 (2013).

Lee, D. J. et al. Septohippocampal Neuromodulation Improves Cognition after Traumatic Brain Injury. Journal of Neurotrauma 32, 1822–1832, doi:10.1089/neu.2014.3744 (2015).

Sweet, J. A., Eakin, K. C., Munyon, C. N. & Miller, J. P. Improved learning and memory with theta-burst stimulation of the fornix in rat model of traumatic brain injury. Hippocampus 24, 1592–1600, doi:10.1002/hipo.22338 (2014).

Chang, J., Phelan, M. & Cummings, B. J. A meta-analysis of efficacy in pre-clinical human stem cell therapies for traumatic brain injury. Experimental Neurology 273, 225–233, doi:10.1016/j.expneurol.2015.08.020 (2015).

Zhu, B., Eom, J. & Hunt, R. F. Transplanted interneurons improve memory precision after traumatic brain injury. Nature Communications 10, 5156, doi:10.1038/s41467-019-13170-w (2019).

Girgis, F., Pace, J., Sweet, J. & Miller, J. P. Hippocampal Neurophysiologic Changes after Mild Traumatic Brain Injury and Potential Neuromodulation Treatment Approaches. Frontiers in Systems Neuroscience 10, 8, doi:10.3389/fnsys.2016.00008 (2016).

Pevzner, A., Izadi, A., Lee, D. J., Shahlaie, K. & Gurkoff, G. G. Making Waves in the Brain: What Are Oscillations, and Why Modulating Them Makes Sense for Brain Injury. Frontiers in Systems Neuroscience 10, 30, doi:10.3389/fnsys.2016.00030 (2016).

Babiloni, C. et al. Brain neural synchronization and functional coupling in Alzheimer’s disease as revealed by resting state EEG rhythms. Int. J. Psychophysiol. 103, 88–102, doi:10.1016/j.ijpsycho.2015.02.008 (2016).

Bazzigaluppi, P. et al. Early-stage attenuation of phase-amplitude coupling in the hippocampus and medial prefrontal cortex in a transgenic rat model of Alzheimer’s disease. J. Neurochem. 144, 669–679, doi:10.1111/jnc.14136 (2018).

Caixeta, F. V., Cornélio, A. M., Scheffer-Teixeira, R., Ribeiro, S. & Tort, A. B. L. Ketamine alters oscillatory coupling in the hippocampus. Sci. Rep. 3, 2348, doi:10.1038/srep02348 (2013).

Chauvière, L. et al. Early Deficits in Spatial Memory and Theta Rhythm in Experimental Temporal Lobe Epilepsy. The Journal of Neuroscience 29, 5402–5410, doi:10.1523/jneurosci.4699-08.2009 (2009).

Dugladze, T. et al. Impaired hippocampal rhythmogenesis in a mouse model of mesial temporal lobe epilepsy. Proceedings of the National Academy of Sciences 104, 17530–17535, doi:10.1073/pnas.0708301104 (2007).

Jacobson, T. K. et al. Hippocampal theta, gamma, and theta-gamma coupling: effects of aging, environmental change, and cholinergic activation. J. Neurophysiol. 109, 1852–1865, doi:10.1152/jn.00409.2012 (2013).

Moran, L. V. & Hong, L. E. High vs Low Frequency Neural Oscillations in Schizophrenia. Schizophr. Bull. 37, 659–663, doi:10.1093/schbul/sbr056 (2011).

Shuman, T. et al. Breakdown of spatial coding and interneuron synchronization in epileptic mice. Nat Neurosci 23, 229–238, doi:10.1038/s41593-019-0559-0 (2020).

Shuman, T., Amendolara, B. & Golshani, P. Theta Rhythmopathy as a Cause of Cognitive Disability in TLE. Epilepsy Curr. 17, 107–111, doi:10.5698/1535-7511.17.2.107 (2017).

Feng, Y. et al. Distinct changes to hippocampal and medial entorhinal circuits emerge across the progression of cognitive deficits in epilepsy. Cell Rep. 44, 115131, doi:10.1016/j.celrep.2024.115131 (2025).

Kundu, B., Brock, A. A., Englot, D. J., Butson, C. R. & Rolston, J. D. Deep brain stimulation for the treatment of disorders of consciousness and cognition in traumatic brain injury patients: a review. Neurosurg. Focus 45, E14, doi:10.3171/2018.5.focus18168 (2018).

Mankin, E. A. & Fried, I. Modulation of Human Memory by Deep Brain Stimulation of the Entorhinal-Hippocampal Circuitry. Neuron 106, 218–235, doi:10.1016/j.neuron.2020.02.024 (2020).

Miller, J. P. et al. Visual-spatial memory may be enhanced with theta burst deep brain stimulation of the fornix: a preliminary investigation with four cases. Brain 138, 1833–1842, doi:10.1093/brain/awv095 (2015).

Ponce, F. A. et al. Bilateral deep brain stimulation of the fornix for Alzheimer’s disease: surgical safety in the ADvance trial. Journal of Neurosurgery 125, 75–84, doi:10.3171/2015.6.jns15716 (2016).

Brignani, D., Manganotti, P., Rossini, P. M. & Miniussi, C. Modulation of cortical oscillatory activity during transcranial magnetic stimulation. Hum. Brain Mapp. 29, 603–612, doi:10.1002/hbm.20423 (2008).

Hallett, M. Transcranial Magnetic Stimulation: A Primer. Neuron 55, 187–199, doi:10.1016/j.neuron.2007.06.026 (2007).

Huerta, P. T. & Volpe, B. T. Transcranial magnetic stimulation, synaptic plasticity and network oscillations. J. Neuroeng. Rehabilitation 6, 7, doi:10.1186/1743-0003-6-7 (2009).

Helfrich, Randolph F. et al. Entrainment of Brain Oscillations by Transcranial Alternating Current Stimulation. Curr. Biol. 24, 333–339, doi:10.1016/j.cub.2013.12.041 (2014).

Vogeti, S., Boetzel, C. & Herrmann, C. S. Entrainment and Spike-Timing Dependent Plasticity – A Review of Proposed Mechanisms of Transcranial Alternating Current Stimulation. Frontiers in Systems Neuroscience 16, 827353, doi:10.3389/fnsys.2022.827353 (2022).

Mueller, J., Legon, W., Opitz, A., Sato, T. F. & Tyler, W. J. Transcranial Focused Ultrasound Modulates Intrinsic and Evoked EEG Dynamics. Brain Stimul. 7, 900–908, doi:10.1016/j.brs.2014.08.008 (2014).

Yuan, Y., Yan, J., Ma, Z. & Li, X. Noninvasive Focused Ultrasound Stimulation Can Modulate Phase-Amplitude Coupling between Neuronal Oscillations in the Rat Hippocampus. Front. Neurosci. 10, 348, doi:10.3389/fnins.2016.00348 (2016).

Pettersen, K. H., Devor, A., Ulbert, I., Dale, A. M. & Einevoll, G. T. Current-source density estimation based on inversion of electrostatic forward solution: Effects of finite extent of neuronal activity and conductivity discontinuities. J Neurosci Meth 154, 116–133, doi:10.1016/j.jneumeth.2005.12.005 (2006).

Berens, P. CircStat : A MATLAB Toolbox for Circular Statistics. J. Stat. Softw. 31, doi:10.18637/jss.v031.i10 (2009).

Arts, L. P. A. & Broek, E. L. v. d. The fast continuous wavelet transformation (fCWT) for real-time, high-quality, noise-resistant time–frequency analysis. Nat. Comput. Sci. 2, 47–58, doi:10.1038/s43588-021-00183-z (2022).

Weber, M. T., Arena, J. D., Xiao, R., Wolf, J. A. & Johnson, V. E. CLARITY reveals a more protracted temporal course of axon swelling and disconnection than previously described following traumatic brain injury. Brain Pathol. 29, 437–450, doi:10.1111/bpa.12677 (2019).

